## Supplementary tables, figures and methods for "Identification of the trail-following pheromone receptor in termites"

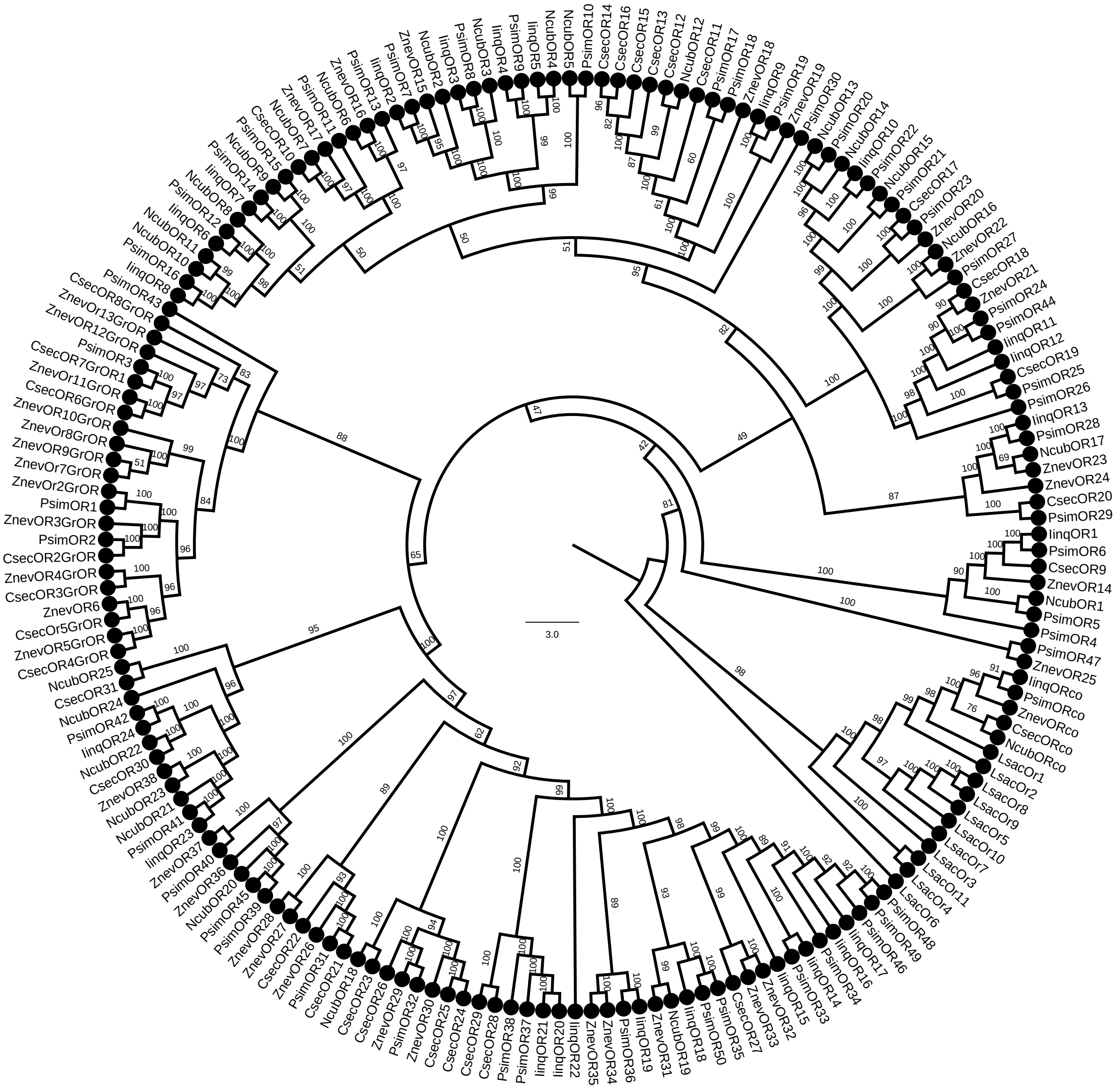

**Fig. S1.** Full version of the phylogenetic tree of termite ORs shown in Fig. 1 of the main text. Protein sequences of termite ORs can be found under the same labeling in Johny et al. (2023). *Lepisma saccharina* sequences used as basal insect outgroup are listed in Thoma et al. (2019). The topology and branching supports were inferred using the IQ-TREE maximum likelihood algorithm with the JTT+F+R8 model and supported by 10,000 iterations of ultrafast bootstrap approximation.

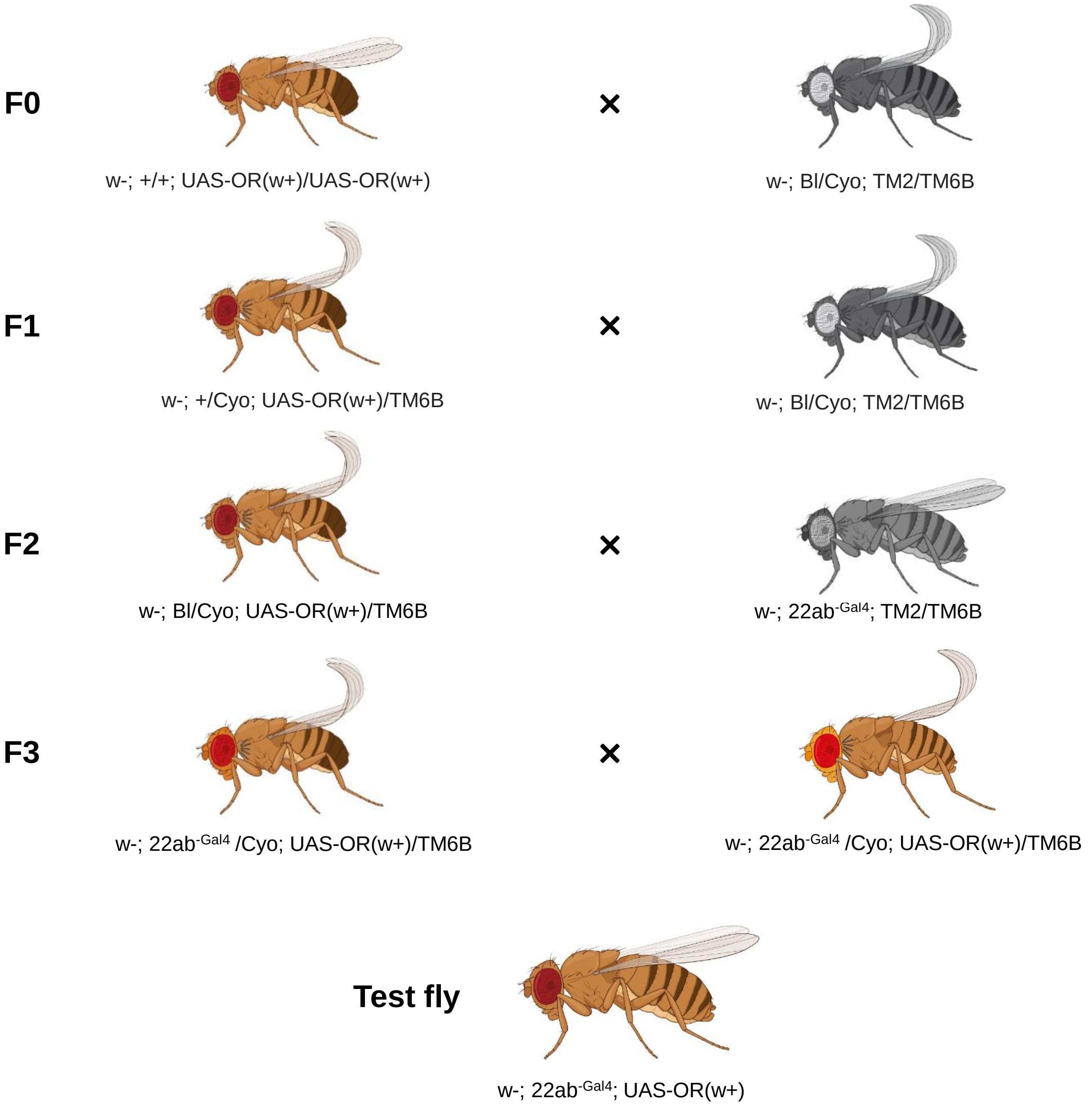


**Fig. S2.** Crossing scheme of termite ORs heterologous expression using *Drosophila melanogaster* empty neurons in ab3 sensilla.

**Table S1.** SSR responses to Panel 1 for PsimOR9. Related to Fig. 2.

| Replicate | Compound | Pre-stimulation spikes | Post-stimulation spikes | Generated spikes | Δ Spikes/s |
|---|---|---|---|---|---|
| 1 | hexane | 8 | 5 | -3 | |
| 1 | (3*Z*,6*Z*)-dodecadien-1-ol | 6 | 9 | 3 | 6 |
| 1 | (3*Z*)-dodecen-1-ol | 11 | 10 | -1 | 2 |
| 1 | (3*Z*,6*Z*,8*E*)-dodecatrien-1-ol | 5 | 12 | 7 | 10 |
| 1 | *n*-eicosane | 4 | 9 | 5 | 8 |
| 1 | 1-octadecanol | 10 | 5 | -5 | -2 |
| 1 | *n*-docosane | 10 | 11 | 1 | 4 |
| 1 | dodec-3-yn-1-ol | 1 | 8 | 7 | 10 |
| 1 | neocembrene | 9 | 11 | 2 | 5 |
| 1 | δ-cadinene | 3 | 0 | -3 | 0 |
| 1 | (3*R*,6*E*)-nerolidol | 0 | 0 | 0 | 3 |
| 2 | hexane | 6 | 3 | -3 | |
| 2 | (3*Z*,6*Z*)-dodecadien-1-ol | 9 | 7 | -2 | 1 |
| 2 | (3*Z*)-dodecen-1-ol | 10 | 8 | -2 | 1 |
| 2 | (3*Z*,6*Z*,8*E*)-dodecatrien-1-ol | 4 | 13 | 9 | 12 |
| 2 | *n*-eicosane | 12 | 2 | -10 | -7 |
| 2 | 1-octadecanol | 8 | 6 | -2 | 1 |
| 2 | *n*-docosane | 11 | 12 | 1 | 4 |
| 2 | dodec-3-yn-1-ol | 9 | 10 | 1 | 4 |
| 2 | neocembrene | 8 | 13 | 5 | 8 |
| 2 | δ-cadinene | 12 | 15 | 3 | 6 |
| 2 | (3*R*,6*E*)-nerolidol | 8 | 14 | 6 | 9 |
| 2 | geranylgeraniol | 12 | 10 | -2 | 1 |
| 3 | hexane | 6 | 5 | -1 | |
| 3 | (3*Z*,6*Z*)-dodecadien-1-ol | 10 | 17 | 7 | 8 |
| 3 | (3*Z*)-dodecen-1-ol | 9 | 14 | 5 | 6 |
| 3 | (3*Z*,6*Z*,8*E*)-dodecatrien-1-ol | 6 | 19 | 13 | 14 |
| 3 | *n*-eicosane | 4 | 15 | 11 | 12 |
| 3 | 1-octadecanol | 13 | 11 | -2 | -1 |
| 3 | *n*-docosane | 7 | 12 | 5 | 6 |
| 3 | dodec-3-yn-1-ol | 8 | 11 | 3 | 4 |
| 3 | neocembrene | 11 | 18 | 7 | 8 |
| 3 | δ-cadinene | 10 | 18 | 8 | 9 |
| 3 | (3*R*,6*E*)-nerolidol | 14 | 11 | -3 | -2 |
| 3 | geranylgeraniol | 7 | 12 | 5 | 6 |
| 4 | hexane | 11 | 10 | -1 | |
| 4 | (3*Z*,6*Z*)-dodecadien-1-ol | 12 | 17 | 5 | 6 |
| 4 | (3*Z*)-dodecen-1-ol | 4 | 15 | 11 | 12 |
| 4 | (3*Z*,6*Z*,8*E*)-dodecatrien-1-ol | 3 | 8 | 5 | 6 |
| 4 | *n*-eicosane | 3 | 3 | 0 | 1 |
| 4 | 1-octadecanol | 2 | 2 | 0 | 1 |
| 4 | *n*-docosane | 5 | 10 | 5 | 6 |
| 4 | dodec-3-yn-1-ol | 4 | 12 | 8 | 9 |
| 4 | neocembrene | 2 | 9 | 7 | 8 |
| 4 | δ-cadinene | 6 | 5 | -1 | 0 |
| 4 | (3*R*,6*E*)-nerolidol | 1 | 4 | 3 | 4 |
| 4 | geranylgeraniol | 3 | 5 | 2 | 3 |
| 5 | hexane | 10 | 8 | -2 | |
| 5 | (3*Z*,6*Z*)-dodecadien-1-ol | 7 | 8 | 1 | 3 |
| 5 | (3*Z*)-dodecen-1-ol | 9 | 9 | 0 | 2 |
| 5 | (3*Z*,6*Z*,8*E*)-dodecatrien-1-ol | 5 | 15 | 10 | 12 |
| 5 | *n*-eicosane | 6 | 12 | 6 | 8 |
| 5 | 1-octadecanol | 9 | 16 | 7 | 9 |
| 5 | *n*-docosane | 8 | 10 | 2 | 4 |
| 5 | dodec-3-yn-1-ol | 6 | 9 | 3 | 5 |
| 5 | neocembrene | 1 | 5 | 4 | 6 |
| 5 | δ-cadinene | 5 | 0 | -5 | -3 |
| 5 | (3*R*,6*E*)-nerolidol | 4 | 2 | -2 | 0 |
| 5 | geranylgeraniol | 3 | 1 | -2 | 0 |

**Table S2.** SSR responses to Panel 1 for PsimOR14. Related to Fig. 2.

| Replicate | Compound | Pre-stimulation spikes | Post-stimulation spikes | Generated spikes | Δ Spikes/s |
|---|---|---|---|---|---|
| 1 | hexane | 7 | 3 | -4 | |
| 1 | 1-octadecanol | 10 | 6 | -4 | 0 |
| 1 | (3*Z*,6*Z*)-dodecadien-1-ol | 6 | 7 | 1 | 5 |
| 1 | (3*Z*)-dodecen-1-ol | 16 | 8 | -8 | -4 |
| 1 | dodec-3-yn-1-ol | 8 | 18 | 10 | 14 |
| 1 | (3*Z*,6*Z*,8*E*)-dodecatrien-1-ol | 13 | 12 | -1 | 3 |
| 1 | (3*R*,6*E*)-nerolidol | 15 | 8 | -7 | -3 |
| 1 | geranylgeraniol | 4 | 22 | 18 | 22 |
| 1 | *n*-docosane | 13 | 10 | -3 | 1 |
| 1 | *n*-eicosane | 10 | 8 | -2 | 2 |
| 1 | neocembrene | 14 | 62 | 48 | 52 |
| 1 | *δ*-cadinene | 13 | 22 | 9 | 13 |
| 2 | hexane | 8 | 9 | 1 | |
| 2 | 1-octadecanol | 5 | 3 | -2 | -3 |
| 2 | (3*Z*,6*Z*)-dodecadien-1-ol | 7 | 3 | -4 | -5 |
| 2 | (3*Z*)-dodecen-1-ol | 13 | 3 | -10 | -11 |
| 2 | dodec-3-yn-1-ol | 9 | 3 | -6 | -7 |
| 2 | (3*Z*,6*Z*,8*E*)-dodecatrien-1-ol | 13 | 3 | -10 | -11 |
| 2 | (3*R*,6*E*)-nerolidol | 6 | 5 | -1 | -2 |
| 2 | geranylgeraniol | 4 | 38 | 34 | 33 |
| 2 | *n*-docosane | 8 | 2 | -6 | -7 |
| 2 | *n*-eicosane | 7 | 2 | -5 | -6 |
| 2 | neocembrene | 4 | 55 | 51 | 50 |
| 2 | *δ*-cadinene | 4 | 5 | 1 | 0 |
| 3 | hexane | 17 | 15 | -2 | |
| 3 | 1-octadecanol | 11 | 6 | -5 | -3 |
| 3 | (3*Z*,6*Z*)-dodecadien-1-ol | 10 | 13 | 3 | 5 |
| 3 | (3*Z*)-dodecen-1-ol | 17 | 7 | -10 | -8 |
| 3 | dodec-3-yn-1-ol | 10 | 7 | -3 | -1 |
| 3 | (3*Z*,6*Z*,8*E*)-dodecatrien-1-ol | 18 | 5 | -13 | -11 |
| 3 | (3*R*,6*E*)-nerolidol | 17 | 14 | -3 | -1 |
| 3 | geranylgeraniol | 15 | 41 | 26 | 28 |
| 3 | *n*-docosane | 20 | 12 | -8 | -6 |
| 3 | *n*-eicosane | 13 | 4 | -9 | -7 |
| 3 | neocembrene | 18 | 90 | 72 | 74 |
| 3 | *δ*-cadinene | 12 | 6 | -6 | -4 |
| 4 | hexane | 6 | 7 | 1 | |
| 4 | 1-octadecanol | 15 | 3 | -12 | -13 |
| 4 | (3*Z*,6*Z*)-dodecadien-1-ol | 7 | 7 | 0 | -1 |
| 4 | (3*Z*)-dodecen-1-ol | 14 | 5 | -9 | -10 |
| 4 | dodec-3-yn-1-ol | 16 | 9 | -7 | -8 |
| 4 | (3*Z*,6*Z*,8*E*)-dodecatrien-1-ol | 12 | 6 | -6 | -7 |
| 4 | (3*R*,6*E*)-nerolidol | 12 | 6 | -6 | -7 |
| 4 | geranylgeraniol | 2 | 28 | 26 | 25 |
| 4 | *n*-docosane | 16 | 4 | -12 | -13 |
| 4 | *n*-eicosane | 7 | 1 | -6 | -7 |
| 4 | neocembrene | 12 | 57 | 45 | 44 |
| 4 | *δ*-cadinene | 5 | 7 | 2 | 1 |
| 5 | hexane | 8 | 6 | -2 | |
| 5 | 1-octadecanol | 8 | 3 | -5 | -3 |
| 5 | (3*Z*,6*Z*)-dodecadien-1-ol | 6 | 3 | -3 | -1 |
| 5 | (3*Z*)-dodecen-1-ol | 2 | 5 | 3 | 5 |
| 5 | dodec-3-yn-1-ol | 2 | 6 | 4 | 6 |
| 5 | (3*Z*,6*Z*,8*E*)-dodecatrien-1-ol | 8 | 4 | -4 | -2 |
| 5 | (3*R*,6*E*)-nerolidol | 10 | 8 | -2 | 0 |
| 5 | geranylgeraniol | 6 | 20 | 14 | 16 |
| 5 | *n*-docosane | 6 | 7 | 1 | 3 |
| 5 | *n*-eicosane | 6 | 4 | -2 | 0 |
| 5 | neocembrene | 5 | 49 | 44 | 46 |
| 5 | *δ*-cadinene | 10 | 7 | -3 | -1 |

**Table S3.** SSR responses to Panel 1 for PsimOR30. Related to Fig. 2.

| Replicate | Compound | Pre-stimulation spikes | Post-stimulation spikes | Generated spikes | Δ Spikes/s |
|---|---|---|---|---|---|
| 1 | hexane | 1 | 0 | -1 | |
| 1 | (3*Z*,6*Z*)-dodecadien-1-ol | 1 | 5 | 4 | 5 |
| 1 | (3*Z*)-dodecen-1-ol | 7 | 3 | -4 | -3 |
| 1 | (3*Z*,6*Z*,8*E*)-dodecatrien-1-ol | 4 | 6 | 2 | 3 |
| 1 | *n*-eicosane | 4 | 8 | 4 | 5 |
| 1 | 1-octadecanol | 5 | 6 | 1 | 2 |
| 1 | *n*-docosane | 5 | 6 | 1 | 2 |
| 1 | dodec-3-yn-1-ol | 4 | 2 | -2 | -1 |
| 1 | neocembrene | 5 | 3 | -2 | -1 |
| 1 | δ-cadinene | 5 | 3 | -2 | -1 |
| 1 | (3*R*,6*E*)-nerolidol | 2 | 3 | 1 | 2 |
| 1 | geranylgeraniol | 4 | 2 | -2 | -1 |
| 2 | hexane | 4 | 4 | 0 | |
| 2 | (3*Z*,6*Z*)-dodecadien-1-ol | 5 | 6 | 1 | 1 |
| 2 | (3*Z*)-dodecen-1-ol | 5 | 8 | 3 | 3 |
| 2 | (3*Z*,6*Z*,8*E*)-dodecatrien-1-ol | 7 | 7 | 0 | 0 |
| 2 | *n*-eicosane | 7 | 7 | 0 | 0 |
| 2 | 1-octadecanol | 9 | 6 | -3 | -3 |
| 2 | *n*-docosane | 5 | 11 | 6 | 6 |
| 2 | dodec-3-yn-1-ol | 9 | 7 | -2 | -2 |
| 2 | neocembrene | 9 | 11 | 2 | 2 |
| 2 | δ-cadinene | 5 | 7 | 2 | 2 |
| 2 | (3*R*,6*E*)-nerolidol | 11 | 8 | -3 | -3 |
| 2 | geranylgeraniol | 10 | 7 | -3 | -3 |
| 3 | hexane | 8 | 5 | -3 | |
| 3 | (3*Z*,6*Z*)-dodecadien-1-ol | 7 | 5 | -2 | 1 |
| 3 | (3*Z*)-dodecen-1-ol | 6 | 5 | -1 | 2 |
| 3 | (3*Z*,6*Z*,8*E*)-dodecatrien-1-ol | 9 | 5 | -4 | -1 |
| 3 | *n*-eicosane | 2 | 10 | 8 | 11 |
| 3 | 1-octadecanol | 7 | 7 | 0 | 3 |
| 3 | *n*-docosane | 8 | 8 | 0 | 3 |
| 3 | dodec-3-yn-1-ol | 3 | 7 | 4 | 7 |
| 3 | neocembrene | 7 | 11 | 4 | 7 |
| 3 | δ-cadinene | 8 | 7 | -1 | 2 |
| 3 | (3*R*,6*E*)-nerolidol | 7 | 3 | -4 | -1 |
| 3 | geranylgeraniol | 5 | 10 | 5 | 8 |

**Table S4.** SSR responses to Panel 1 for PsimOR31. Related to Fig. 2.

| Replicate | Compound | Pre-stimulation spikes | Post-stimulation spikes | Generated spikes | Δ Spikes/s |
|---|---|---|---|---|---|
| 1 | hexane | 5 | 6 | 1 | |
| 1 | (3*Z*,6*Z*)-dodecadien-1-ol | 5 | 24 | 19 | 18 |
| 1 | (3*Z*)-dodecen-1-ol | 5 | 25 | 20 | 19 |
| 1 | (3*Z*,6*Z*,8*E*)-dodecatrien-1-ol | 3 | 27 | 24 | 23 |
| 1 | *n*-eicosane | 3 | 25 | 22 | 21 |
| 1 | 1-octadecanol | 5 | 25 | 20 | 19 |
| 1 | *n*-docosane | 6 | 27 | 21 | 20 |
| 1 | dodec-3-yn-1-ol | 4 | 28 | 24 | 23 |
| 1 | neocembrene | 6 | 19 | 13 | 12 |
| 1 | *δ*-cadinene | 3 | 19 | 16 | 15 |
| 1 | (3*R*,6*E*)-nerolidol | 5 | 16 | 11 | 10 |
| 1 | geranylgeraniol | 3 | 10 | 7 | 6 |
| 2 | hexane | 3 | 2 | -1 | |
| 2 | (3*Z*,6*Z*)-dodecadien-1-ol | 5 | 5 | 0 | 1 |
| 2 | (3*Z*)-dodecen-1-ol | 4 | 9 | 5 | 6 |
| 2 | (3*Z*,6*Z*,8*E*)-dodecatrien-1-ol | 2 | 7 | 5 | 6 |
| 2 | *n*-eicosane | 5 | 10 | 5 | 6 |
| 2 | 1-octadecanol | 6 | 6 | 0 | 1 |
| 2 | *n*-docosane | 5 | 4 | -1 | 0 |
| 2 | dodec-3-yn-1-ol | 7 | 6 | -1 | 0 |
| 2 | neocembrene | 2 | 5 | 3 | 4 |
| 2 | *δ*-cadinene | 2 | 10 | 8 | 9 |
| 2 | (3*R*,6*E*)-nerolidol | 2 | 2 | 0 | 1 |
| 3 | hexane | 4 | 3 | -1 | |
| 3 | (3*Z*,6*Z*)-dodecadien-1-ol | 9 | 7 | -2 | -1 |
| 3 | (3*Z*)-dodecen-1-ol | 3 | 14 | 11 | 12 |
| 3 | (3*Z*,6*Z*,8*E*)-dodecatrien-1-ol | 8 | 14 | 6 | 7 |
| 3 | *n*-eicosane | 9 | 21 | 12 | 13 |
| 3 | 1-octadecanol | 6 | 21 | 15 | 16 |
| 3 | *n*-docosane | 6 | 27 | 21 | 22 |
| 3 | dodec-3-yn-1-ol | 10 | 18 | 8 | 9 |
| 3 | neocembrene | 9 | 17 | 8 | 9 |
| 3 | *δ*-cadinene | 7 | 20 | 13 | 14 |
| 3 | (3*R*,6*E*)-nerolidol | 6 | 14 | 8 | 9 |
| 3 | geranylgeraniol | 10 | 21 | 11 | 12 |
| 4 | hexane | 13 | 15 | 2 | |
| 4 | (3*Z*,6*Z*)-dodecadien-1-ol | 20 | 17 | -3 | -5 |
| 4 | (3*Z*)-dodecen-1-ol | 15 | 19 | 4 | 2 |
| 4 | (3*Z*,6*Z*,8*E*)-dodecatrien-1-ol | 14 | 21 | 7 | 5 |
| 4 | *n*-eicosane | 18 | 22 | 4 | 2 |
| 4 | 1-octadecanol | 19 | 22 | 3 | 1 |
| 4 | *n*-docosane | 16 | 20 | 4 | 2 |
| 4 | dodec-3-yn-1-ol | 11 | 23 | 12 | 10 |
| 4 | neocembrene | 8 | 22 | 14 | 12 |
| 4 | *δ*-cadinene | 10 | 24 | 14 | 12 |
| 4 | (3*R*,6*E*)-nerolidol | 16 | 21 | 5 | 3 |
| 4 | geranylgeraniol | 12 | 21 | 9 | 7 |
| 5 | hexane | 11 | 10 | -1 | |
| 5 | (3*Z*,6*Z*)-dodecadien-1-ol | 17 | 18 | 1 | 2 |
| 5 | (3*Z*)-dodecen-1-ol | 5 | 16 | 11 | 12 |
| 5 | (3*Z*,6*Z*,8*E*)-dodecatrien-1-ol | 8 | 12 | 4 | 5 |
| 5 | *n*-eicosane | 17 | 14 | -3 | -2 |
| 5 | 1-octadecanol | 7 | 13 | 6 | 7 |
| 5 | *n*-docosane | 11 | 16 | 5 | 6 |
| 5 | dodec-3-yn-1-ol | 10 | 15 | 5 | 6 |
| 5 | neocembrene | 0 | 15 | 15 | 16 |
| 6 | hexane | 24 | 25 | 1 | |
| 6 | (3*Z*,6*Z*)-dodecadien-1-ol | 21 | 46 | 25 | 24 |
| 6 | (3*Z*)-dodecen-1-ol | 18 | 44 | 26 | 25 |
| 6 | (3*Z*,6*Z*,8*E*)-dodecatrien-1-ol | 14 | 44 | 30 | 29 |
| 6 | *n*-eicosane | 17 | 43 | 26 | 25 |
| 6 | 1-octadecanol | 10 | 44 | 34 | 33 |
| 6 | *n*-docosane | 12 | 38 | 26 | 25 |
| 6 | dodec-3-yn-1-ol | 17 | 47 | 30 | 29 |
| 6 | neocembrene | 11 | 32 | 21 | 20 |
| 6 | *δ*-cadinene | 19 | 27 | 8 | 7 |
| 6 | (3*R*,6*E*)-nerolidol | 7 | 26 | 19 | 18 |
| 6 | geranylgeraniol | 9 | 19 | 10 | 9 |

**Table S5.** SSR responses to Panel 1 for PsimOR14 vs. $W^{1118}$. Related to Fig. 2.

| Fly line | Replicate | Compound | Pre-stimulation spikes | Post-stimulation spikes | Generated spikes | Δ Spikes/s |
|---|---|---|---|---|---|---|
| PsimOR14 | 1 | hexane | 7 | 3 | -4 | |
| PsimOR14 | 1 | (3*Z*,6*Z*)-dodecadien-1-ol | 6 | 7 | 1 | 5 |
| PsimOR14 | 1 | (3*Z*)-dodecen-1-ol | 16 | 8 | -8 | -4 |
| PsimOR14 | 1 | (3*Z*,6*Z*,8*E*)-dodecatrien-1-ol | 13 | 12 | -1 | 3 |
| PsimOR14 | 1 | *n*-eicosane | 10 | 8 | -2 | 2 |
| PsimOR14 | 1 | 1-octadecanol | 10 | 6 | -4 | 0 |
| PsimOR14 | 1 | *n*-docosane | 13 | 10 | -3 | 1 |
| PsimOR14 | 1 | dodec-3-yn-1-ol | 8 | 18 | 10 | 14 |
| PsimOR14 | 1 | neocembrene | 14 | 62 | 48 | 52 |
| PsimOR14 | 1 | *δ*-cadinene | 13 | 22 | 9 | 13 |
| PsimOR14 | 1 | (3*R*,6*E*)-nerolidol | 15 | 8 | -7 | -3 |
| PsimOR14 | 1 | geranylgeraniol | 4 | 22 | 18 | 22 |
| PsimOR14 | 2 | hexane | 8 | 9 | 1 | |
| PsimOR14 | 2 | (3*Z*,6*Z*)-dodecadien-1-ol | 7 | 3 | -4 | -5 |
| PsimOR14 | 2 | (3*Z*)-dodecen-1-ol | 13 | 3 | -10 | -11 |
| PsimOR14 | 2 | (3*Z*,6*Z*,8*E*)-dodecatrien-1-ol | 13 | 3 | -10 | -11 |
| PsimOR14 | 2 | *n*-eicosane | 7 | 2 | -5 | -6 |
| PsimOR14 | 2 | 1-octadecanol | 5 | 3 | -2 | -3 |
| PsimOR14 | 2 | *n*-docosane | 8 | 2 | -6 | -7 |
| PsimOR14 | 2 | dodec-3-yn-1-ol | 9 | 3 | -6 | -7 |
| PsimOR14 | 2 | neocembrene | 4 | 55 | 51 | 50 |
| PsimOR14 | 2 | *δ*-cadinene | 4 | 5 | 1 | 0 |
| PsimOR14 | 2 | (3*R*,6*E*)-nerolidol | 6 | 5 | -1 | -2 |
| PsimOR14 | 2 | geranylgeraniol | 4 | 38 | 34 | 33 |
| PsimOR14 | 3 | hexane | 17 | 15 | -2 | |
| PsimOR14 | 3 | (3*Z*,6*Z*)-dodecadien-1-ol | 10 | 13 | 3 | 5 |
| PsimOR14 | 3 | (3*Z*)-dodecen-1-ol | 17 | 7 | -10 | -8 |
| PsimOR14 | 3 | (3*Z*,6*Z*,8*E*)-dodecatrien-1-ol | 18 | 5 | -13 | -11 |
| PsimOR14 | 3 | *n*-eicosane | 13 | 4 | -9 | -7 |
| PsimOR14 | 3 | 1-octadecanol | 11 | 6 | -5 | -3 |
| PsimOR14 | 3 | *n*-docosane | 20 | 12 | -8 | -6 |
| PsimOR14 | 3 | dodec-3-yn-1-ol | 10 | 7 | -3 | -1 |
| PsimOR14 | 3 | neocembrene | 18 | 90 | 72 | 74 |
| PsimOR14 | 3 | *δ*-cadinene | 12 | 6 | -6 | -4 |
| PsimOR14 | 3 | (3*R*,6*E*)-nerolidol | 17 | 14 | -3 | -1 |
| PsimOR14 | 3 | geranylgeraniol | 15 | 41 | 26 | 28 |
| PsimOR14 | 4 | hexane | 6 | 7 | 1 | |
| PsimOR14 | 4 | neocembrene | 12 | 57 | 45 | 44 |
| PsimOR14 | 4 | geranylgeraniol | 2 | 28 | 26 | 25 |
| PsimOR14 | 4 | (3*Z*,6*Z*)-dodecadien-1-ol | 7 | 7 | 0 | -1 |
| PsimOR14 | 4 | (3*Z*)-dodecen-1-ol | 14 | 5 | -9 | -10 |
| PsimOR14 | 4 | (3*Z*,6*Z*,8*E*)-dodecatrien-1-ol | 12 | 6 | -6 | -7 |
| PsimOR14 | 4 | *n*-eicosane | 7 | 1 | -6 | -7 |
| PsimOR14 | 4 | 1-octadecanol | 15 | 3 | -12 | -13 |
| PsimOR14 | 4 | *n*-docosane | 16 | 4 | -12 | -13 |
| PsimOR14 | 4 | dodec-3-yn-1-ol | 16 | 9 | -7 | -8 |
| PsimOR14 | 4 | *δ*-cadinene | 5 | 7 | 2 | 1 |
| PsimOR14 | 4 | (3*R*,6*E*)-nerolidol | 12 | 6 | -6 | -7 |
| PsimOR14 | 5 | hexane | 8 | 6 | -2 | |
| PsimOR14 | 5 | neocembrene | 5 | 49 | 44 | 46 |
| PsimOR14 | 5 | geranylgeraniol | 6 | 20 | 14 | 16 |
| PsimOR14 | 5 | (3*Z*,6*Z*)-dodecadien-1-ol | 6 | 3 | -3 | -1 |
| PsimOR14 | 5 | (3*Z*)-dodecen-1-ol | 2 | 5 | 3 | 5 |
| PsimOR14 | 5 | (3*Z*,6*Z*,8*E*)-dodecatrien-1-ol | 8 | 4 | -4 | -2 |
| PsimOR14 | 5 | *n*-eicosane | 6 | 4 | -2 | 0 |
| PsimOR14 | 5 | 1-octadecanol | 8 | 3 | -5 | -3 |
| PsimOR14 | 5 | *n*-docosane | 6 | 7 | 1 | 3 |
| PsimOR14 | 5 | dodec-3-yn-1-ol | 2 | 6 | 4 | 6 |
| PsimOR14 | 5 | *δ*-cadinene | 10 | 7 | -3 | -1 |
| PsimOR14 | 5 | (3*R*,6*E*)-nerolidol | 10 | 8 | -2 | 0 |

| Fly line | Replicate | Compound | Pre-stimulation spikes | Post-stimulation spikes | Generated spikes | Δ Spikes/s |
|---|---|---|---|---|---|---|
| W1118 | 1 | hexane | 3 | 6 | 3 | |
| W1118 | 1 | (3*Z*,6*Z*)-dodecadien-1-ol | 1 | 3 | 2 | -1 |
| W1118 | 1 | (3*Z*)-dodecen-1-ol | 2 | 4 | 2 | -1 |
| W1118 | 1 | (3*Z*,6*Z*,8*E*)-dodecatrien-1-ol | 4 | 1 | -3 | -6 |
| W1118 | 1 | *n*-eicosane | 1 | 19 | 18 | 15 |
| W1118 | 1 | 1-octadecanol | 4 | 2 | -2 | -5 |
| W1118 | 1 | *n*-docosane | 2 | 6 | 4 | 1 |
| W1118 | 1 | dodec-3-yn-1-ol | 2 | 5 | 3 | 0 |
| W1118 | 1 | neocembrene | 2 | 2 | 0 | -3 |
| W1118 | 1 | *δ*-cadinene | 1 | 4 | 3 | 0 |
| W1118 | 1 | (3*R*,6*E*)-nerolidol | 3 | 1 | -2 | -5 |
| W1118 | 1 | geranylgeraniol | 3 | 4 | 1 | -2 |
| W1118 | 2 | hexane | 1 | 4 | 3 | |
| W1118 | 2 | (3*Z*,6*Z*)-dodecadien-1-ol | 2 | 4 | 2 | -1 |
| W1118 | 2 | (3*Z*)-dodecen-1-ol | 5 | 4 | -1 | -4 |
| W1118 | 2 | (3*Z*,6*Z*,8*E*)-dodecatrien-1-ol | 5 | 3 | -2 | -5 |
| W1118 | 2 | *n*-eicosane | 3 | 3 | 0 | -3 |
| W1118 | 2 | 1-octadecanol | 3 | 2 | -1 | -4 |
| W1118 | 2 | *n*-docosane | 0 | 5 | 5 | 2 |
| W1118 | 2 | dodec-3-yn-1-ol | 3 | 3 | 0 | -3 |
| W1118 | 2 | neocembrene | 4 | 4 | 0 | -3 |
| W1118 | 2 | *δ*-cadinene | 6 | 0 | -6 | -9 |
| W1118 | 2 | (3*R*,6*E*)-nerolidol | 1 | 3 | 2 | -1 |
| W1118 | 2 | geranylgeraniol | 3 | 4 | 1 | -2 |
| W1118 | 3 | hexane | 5 | 1 | -4 | |
| W1118 | 3 | (3Z,6Z)-dodecadien-1-ol | 5 | 5 | 0 | 4 |
| W1118 | 3 | (3Z)-dodecen-1-ol | 5 | 2 | -3 | 1 |
| W1118 | 3 | (3Z,6Z,8E)-dodecatrien-1-ol | 2 | 4 | 2 | 6 |
| W1118 | 3 | n-eicosane | 3 | 5 | 2 | 6 |
| W1118 | 3 | 1-octadecanol | 3 | 5 | 2 | 6 |
| W1118 | 3 | n-docosane | 4 | 5 | 1 | 5 |
| W1118 | 3 | dodec-3-yn-1-ol | 6 | 5 | -1 | 3 |
| W1118 | 3 | neocembrene | 1 | 5 | 4 | 8 |
| W1118 | 3 | δ-cadinene | 4 | 4 | 0 | 4 |
| W1118 | 3 | (3*R*,6*E*)-nerolidol | 6 | 3 | -3 | 1 |
| W1118 | 3 | geranylgeraniol | 4 | 2 | -2 | 2 |
| W1118 | 4 | hexane | 4 | 4 | 0 | |
| W1118 | 4 | (3Z,6Z)-dodecadien-1-ol | 1 | 6 | 5 | 5 |
| W1118 | 4 | (3Z)-dodecen-1-ol | 3 | 3 | 0 | 0 |
| W1118 | 4 | (3Z,6Z,8E)-dodecatrien-1-ol | 6 | 4 | -2 | -2 |
| W1118 | 4 | n-eicosane | 4 | 7 | 3 | 3 |
| W1118 | 4 | 1-octadecanol | 5 | 7 | 2 | 2 |
| W1118 | 4 | n-docosane | 4 | 9 | 5 | 5 |
| W1118 | 4 | dodec-3-yn-1-ol | 0 | 4 | 4 | 4 |
| W1118 | 4 | neocembrene | 2 | 0 | -2 | -2 |
| W1118 | 4 | δ-cadinene | 5 | 2 | -3 | -3 |
| W1118 | 4 | (3*R*,6*E*)-nerolidol | 4 | 1 | -3 | -3 |
| W1118 | 4 | geranylgeraniol | 2 | 0 | -2 | -2 |
| W1118 | 5 | hexane | 3 | 8 | 5 | |
| W1118 | 5 | (3Z,6Z)-dodecadien-1-ol | 6 | 11 | 5 | 0 |
| W1118 | 5 | (3Z)-dodecen-1-ol | 9 | 7 | -2 | -7 |
| W1118 | 5 | (3Z,6Z,8E)-dodecatrien-1-ol | 6 | 9 | 3 | -2 |
| W1118 | 5 | n-eicosane | 7 | 14 | 7 | 2 |
| W1118 | 5 | 1-octadecanol | 4 | 13 | 9 | 4 |
| W1118 | 5 | n-docosane | 8 | 7 | -1 | -6 |
| W1118 | 5 | dodec-3-yn-1-ol | 7 | 8 | 1 | -4 |
| W1118 | 5 | neocembrene | 6 | 8 | 2 | -3 |
| W1118 | 5 | δ-cadinene | 5 | 8 | 3 | -2 |
| W1118 | 5 | (3*R*,6*E*)-nerolidol | 6 | 6 | 0 | -5 |
| W1118 | 5 | geranylgeraniol | 6 | 6 | 0 | -5 |

**Table S6.** SSR dose response data for neocembrene and PsimOR14 fly line. Related to Fig. 2.

| | replicate | 1 | 2 | 3 | 4 | 5 | 6 | 7 | 8 | 9 |
|---|---|---|---|---|---|---|---|---|---|---|
| Dose | 0.01 ng | 22 | 10 | 10 | 6 | -4 | 10 | 18 | 16 | 24 |
| | 0.1 ng | 4 | 26 | 26 | 8 | 0 | 0 | 4 | 38 | 12 |
| | 1 ng | 14 | 32 | 14 | 10 | 12 | 2 | 22 | 28 | 10 |
| | 10 ng | 10 | 30 | 32 | 26 | 26 | 14 | 20 | 32 | 6 |
| | 100 ng | 30 | | 24 | 26 | 42 | 38 | 46 | 26 | 10 |
| | 500 ng | 44 | | 36 | 22 | 34 | | | | |

**Table S7.** SSR responses to Panel 2 for PsimOR14. Related to Fig. 3.

| Panel | Replicate | Compound | Pre-stimulation spikes | Post-stimulation spikes | Generated spikes | Δ Spikes/s |
|---|---|---|---|---|---|---|
| 2 | 1 | paraffin oil | 6 | 2 | -4 | |
| 2 | 1 | myrcene | 10 | 15 | 5 | 9 |
| 2 | 1 | eucalyptol | 2 | 2 | 0 | 4 |
| 2 | 1 | (+)-3-carene | 2 | 5 | 3 | 7 |
| 2 | 1 | ethanol | 2 | 6 | 4 | 8 |
| 2 | 1 | (2*E*)-hexenal | 3 | 5 | 2 | 6 |
| 2 | 1 | (+)-*α*-pinene | 3 | 2 | -1 | 3 |
| 2 | 1 | *p*-cymene | 3 | 2 | -1 | 3 |
| 2 | 1 | (+)-limonene | 4 | 3 | -1 | 3 |
| 2 | 1 | (+)-longifolene | 3 | 2 | -1 | 3 |
| 2 | 1 | (-)-*β*-caryophyllene | 1 | 2 | 1 | 5 |
| 2 | 1 | (2*E*)-hexen-1-ol | 1 | 0 | -1 | 3 |
| 2 | 1 | isoamyl propionate | 0 | 0 | 0 | 4 |
| 2 | 1 | sabinene | 2 | 1 | -1 | 3 |
| 2 | 1 | *γ*-terpinene | 0 | 2 | 2 | 6 |
| 2 | 1 | terpinolene | 2 | 0 | -2 | 2 |
| 2 | 1 | toluene | 1 | 5 | 4 | 8 |
| 2 | 1 | pentyl acetate | 1 | 3 | 2 | 6 |
| 2 | 1 | ethyl acetate | 2 | 1 | -1 | 3 |
| 2 | 1 | heptanal | 0 | 0 | 0 | 4 |
| 2 | 1 | *γ*-nonalactone | 2 | 0 | -2 | 2 |
| 2 | 2 | paraffin oil | 0 | 1 | 1 | |
| 2 | 2 | myrcene | 6 | 11 | 5 | 4 |
| 2 | 2 | eucalyptol | 0 | 13 | 13 | 12 |
| 2 | 2 | (+)-3-carene | 6 | 10 | 4 | 3 |
| 2 | 2 | ethanol | 2 | 12 | 10 | 9 |
| 2 | 2 | (2*E*)-hexenal | 6 | 11 | 5 | 4 |
| 2 | 2 | (+)-*α*-pinene | 0 | 10 | 10 | 9 |
| 2 | 2 | *p*-cymene | 8 | 12 | 4 | 3 |
| 2 | 2 | (+)-limonene | 2 | 11 | 9 | 8 |
| 2 | 2 | (+)-longifolene | 1 | 10 | 9 | 8 |
| 2 | 2 | (-)-*β*-caryophyllene | 5 | 7 | 2 | 1 |
| 2 | 2 | (2*E*)-hexen-1-ol | 8 | 12 | 4 | 3 |
| 2 | 2 | isoamyl propionate | 8 | 8 | 0 | -1 |
| 2 | 2 | sabinene | 6 | 16 | 10 | 9 |
| 2 | 2 | *γ*-terpinene | 9 | 7 | -2 | -3 |
| 2 | 2 | terpinolene | 5 | 10 | 5 | 4 |
| 2 | 2 | toluene | 8 | 8 | 0 | -1 |
| 2 | 2 | pentyl acetate | 7 | 9 | 2 | 1 |
| 2 | 2 | ethyl acetate | 8 | 9 | 1 | 0 |
| 2 | 2 | heptanal | 10 | 9 | -1 | -2 |
| 2 | 2 | *γ*-nonalactone | 6 | 12 | 6 | 5 |
| 2 | 3 | paraffin oil | 5 | 6 | 1 | |
| 2 | 3 | myrcene | 2 | 6 | 4 | 3 |
| 2 | 3 | eucalyptol | 2 | 10 | 8 | 7 |
| 2 | 3 | (+)-3-carene | 0 | 14 | 14 | 13 |
| 2 | 3 | ethanol | 2 | 6 | 4 | 3 |
| 2 | 3 | (2*E*)-hexenal | 2 | 12 | 10 | 9 |
| 2 | 3 | (+)-*α*-pinene | 2 | 9 | 7 | 6 |
| 2 | 3 | *p*-cymene | 3 | 4 | 1 | 0 |
| 2 | 3 | (+)-limonene | 1 | 1 | 0 | -1 |
| 2 | 3 | (+)-longifolene | 0 | 10 | 10 | 9 |
| 2 | 3 | (-)-*β*-caryophyllene | 0 | 10 | 10 | 9 |
| 2 | 3 | (2*E*)-hexen-1-ol | 4 | 1 | -3 | -4 |
| 2 | 3 | (2*E*)-hexen-1-ol | 3 | 1 | -2 | -3 |
| 2 | 3 | sabinene | 1 | 2 | 1 | 0 |
| 2 | 3 | *γ*-terpinene | 9 | 5 | -4 | -5 |
| 2 | 3 | terpinolene | 1 | 9 | 8 | 7 |
| 2 | 3 | toluene | 2 | 0 | -2 | -3 |
| 2 | 3 | pentyl acetate | 1 | 9 | 8 | 7 |
| 2 | 3 | ethyl acetate | 4 | 0 | -4 | -5 |
| 2 | 3 | heptanal | 1 | 4 | 3 | 2 |
| 2 | 3 | *γ*-nonalactone | 2 | 2 | 0 | -1 |
| 2 | 4 | paraffin oil | 4 | 5 | 1 | 0 |
| 2 | 4 | myrcene | 3 | 1 | -2 | -3 |
| 2 | 4 | eucalyptol | 0 | 7 | 7 | 6 |
| 2 | 4 | (+)-3-carene | 2 | 5 | 3 | 2 |
| 2 | 4 | ethanol | 1 | 4 | 3 | 2 |
| 2 | 4 | (2*E*)-hexenal | 0 | 3 | 3 | 2 |
| 2 | 4 | (+)-*α*-pinene | 3 | 12 | 9 | 8 |
| 2 | 4 | *p*-cymene | 1 | 11 | 10 | 9 |
| 2 | 4 | (+)-limonene | 0 | 3 | 3 | 2 |
| 2 | 4 | (+)-longifolene | 0 | 8 | 8 | 7 |
| 2 | 4 | (-)-*β*-caryophyllene | 0 | 2 | 2 | 1 |
| 2 | 4 | (2*E*)-hexen-1-ol | 0 | 5 | 5 | 4 |
| 2 | 4 | isoamyl propionate | 0 | 5 | 5 | 4 |
| 2 | 4 | sabinene | 2 | 2 | 0 | -1 |
| 2 | 4 | *γ*-terpinene | 3 | 9 | 6 | 5 |
| 2 | 4 | terpinolene | 7 | 4 | -3 | -4 |
| 2 | 4 | toluene | 2 | 2 | 0 | -1 |
| 2 | 4 | pentyl acetate | 3 | 2 | -1 | -2 |
| 2 | 4 | ethyl acetate | 1 | 1 | 0 | -1 |
| 2 | 4 | heptanal | 3 | 1 | -2 | -3 |
| 2 | 4 | paraffin oil | 6 | 7 | 1 | 0 |
| 2 | 5 | myrcene | 11 | 11 | 0 | -1 |
| 2 | 5 | eucalyptol | 7 | 13 | 6 | 5 |
| 2 | 5 | (+)-3-carene | 12 | 11 | -1 | -2 |
| 2 | 5 | ethanol | 11 | 9 | -2 | -3 |
| 2 | 5 | (2*E*)-hexenal | 11 | 13 | 2 | 1 |
| 2 | 5 | (+)-*α*-pinene | 12 | 11 | -1 | -2 |
| 2 | 5 | *p*-cymene | 11 | 16 | 5 | 4 |
| 2 | 5 | (+)-limonene | 9 | 9 | 0 | -1 |
| 2 | 5 | (+)-longifolene | 17 | 9 | -8 | -9 |
| 2 | 5 | (-)-*β*-caryophyllene | 6 | 13 | 7 | 6 |
| 2 | 5 | (2*E*)-hexen-1-ol | 9 | 3 | -6 | -7 |
| 2 | 5 | isoamyl propionate | 0 | 0 | 0 | -1 |
| 2 | 5 | sabinene | 6 | 19 | 13 | 12 |
| 2 | 5 | *γ*-terpinene | 3 | 13 | 10 | 9 |
| 2 | 5 | terpinolene | 9 | 15 | 6 | 5 |
| 2 | 5 | toluene | 10 | 13 | 3 | 2 |
| 2 | 5 | pentyl acetate | 6 | 11 | 5 | 4 |
| 2 | 5 | ethyl acetate | 8 | 19 | 11 | 10 |
| 2 | 5 | heptanal | 14 | 7 | -7 | -8 |
| 2 | 5 | *γ*-nonalactone | 10 | 14 | 4 | 3 |

**Table S8.** SSR responses to Panel 3 for PsimOR14. Related to Fig. 3.

| Panel | Replicate | Compound | Pre-stimulation spikes | Post-stimulation spikes | Generated spikes | Δ Spikes/s |
|---|---|---|---|---|---|---|
| 3 | 1 | paraffin oil | 4 | 5 | 1 | |
| 3 | 1 | ethyl (2*E*,4*Z*)-decadienoate | 4 | 6 | 2 | 1 |
| 3 | 1 | *p*-cresol | 2 | 9 | 7 | 6 |
| 3 | 1 | *β*-bisabolene | 7 | 2 | -5 | -6 |
| 3 | 1 | (*E*)-myrcenol | 1 | 6 | 5 | 4 |
| 3 | 1 | nonan-1-ol | 1 | 8 | 7 | 6 |
| 3 | 1 | isoamyl acetate | 5 | 11 | 6 | 5 |
| 3 | 1 | 2-phenylethyl acetate | 3 | 9 | 6 | 5 |
| 3 | 1 | octan-1-ol | 6 | 12 | 6 | 5 |
| 3 | 1 | benzyl alcohol | 12 | 11 | -1 | -2 |
| 3 | 1 | methyl jasmonate | 8 | 13 | 5 | 4 |
| 3 | 1 | (*E*)-*β*-ocimene | 6 | 14 | 8 | 7 |
| 3 | 1 | eucarvone | 10 | 4 | -6 | -7 |
| 3 | 1 | *α*-camphorene | 2 | 14 | 12 | 11 |
| 3 | 1 | (-)-trans-pinocarveol | 6 | 4 | -2 | -3 |
| 3 | 1 | cryptone | 8 | 8 | 0 | -1 |
| 3 | 1 | (+)-cis-carveol | 12 | 17 | 5 | 4 |
| 3 | 2 | paraffin oil | 9 | 5 | -4 | |
| 3 | 2 | ethyl (2*E*,4*Z*)-decadienoate | 19 | 11 | -8 | -4 |
| 3 | 2 | *p*-cresol | 15 | 15 | 0 | 4 |
| 3 | 2 | *β*-bisabolene | 8 | 14 | 6 | 10 |
| 3 | 2 | (*E*)-myrcenol | 17 | 16 | -1 | 3 |
| 3 | 2 | nonan-1-ol | 17 | 14 | -3 | 1 |
| 3 | 2 | isoamyl acetate | 20 | 16 | -4 | 0 |
| 3 | 2 | 2-phenylethyl acetate | 13 | 7 | -6 | -2 |
| 3 | 2 | octan-1-ol | 10 | 3 | -7 | -3 |
| 3 | 2 | benzyl alcohol | 3 | 2 | -1 | 3 |
| 3 | 2 | methyl jasmonate | 0 | 1 | 1 | 5 |
| 3 | 2 | (*E*)-*β*-ocimene | 0 | 2 | 2 | 6 |
| 3 | 2 | eucarvone | 0 | 1 | 1 | 5 |
| 3 | 2 | *α*-camphorene | 0 | 1 | 1 | 5 |
| 3 | 2 | (-)-trans-pinocarveol | 0 | 1 | 1 | 5 |
| 3 | 2 | cryptone | 0 | 0 | 0 | 4 |
| 3 | 2 | (+)-cis-carveol | 0 | 0 | 0 | 4 |
| 3 | 3 | paraffin oil | 0 | 1 | 1 | |
| 3 | 3 | ethyl (2*E*,4*Z*)-decadienoate | 4 | 8 | 4 | 3 |
| 3 | 3 | *p*-cresol | 2 | 6 | 4 | 3 |
| 3 | 3 | *β*-bisabolene | 12 | 3 | -9 | -10 |
| 3 | 3 | (*E*)-myrcenol | 6 | 10 | 4 | 3 |
| 3 | 3 | nonan-1-ol | 5 | 7 | 2 | 1 |
| 3 | 3 | isoamyl acetate | 4 | 9 | 5 | 4 |
| 3 | 3 | 2-phenylethyl acetate | 6 | 9 | 3 | 2 |
| 3 | 3 | octan-1-ol | 0 | 14 | 14 | 13 |
| 3 | 3 | benzyl alcohol | 8 | 7 | -1 | -2 |
| 3 | 3 | methyl jasmonate | 9 | 11 | 2 | 1 |
| 3 | 3 | (*E*)-*β*-ocimene | 2 | 4 | 2 | 1 |
| 3 | 3 | eucarvone | 10 | 1 | -9 | -10 |
| 3 | 3 | *α*-camphorene | 7 | 7 | 0 | -1 |
| 3 | 3 | (-)-trans-pinocarveol | 5 | 0 | -5 | -6 |
| 3 | 3 | cryptone | 5 | 8 | 3 | 2 |
| 3 | 3 | (+)-cis-carveol | 5 | 6 | 1 | 0 |
| 3 | 4 | paraffin oil | 2 | 7 | 5 | |
| 3 | 4 | ethyl (2*E*,4*Z*)-decadienoate | 1 | 0 | -1 | -6 |
| 3 | 4 | *p*-cresol | 6 | 10 | 4 | -1 |
| 3 | 4 | *β*-bisabolene | 3 | 9 | 6 | 1 |
| 3 | 4 | (*E*)-myrcenol | 2 | 9 | 7 | 2 |
| 3 | 4 | nonan-1-ol | 10 | 4 | -6 | -11 |
| 3 | 4 | isoamyl acetate | 1 | 6 | 5 | 0 |
| 3 | 4 | 2-phenylethyl acetate | 0 | 0 | 0 | -5 |
| 3 | 4 | octan-1-ol | 2 | 0 | -2 | -7 |
| 3 | 4 | benzyl alcohol | 1 | 5 | 4 | -1 |
| 3 | 4 | methyl jasmonate | 10 | 3 | -7 | -12 |
| 3 | 4 | (*E*)-*β*-ocimene | 0 | 5 | 5 | 0 |
| 3 | 4 | eucarvone | 3 | 1 | -2 | -7 |
| 3 | 4 | *α*-camphorene | 1 | 0 | -1 | -6 |
| 3 | 4 | (-)-trans-pinocarveol | 9 | 5 | -4 | -9 |
| 3 | 4 | cryptone | 9 | 6 | -3 | -8 |
| 3 | 4 | (+)-cis-carveol | 1 | 0 | -1 | -6 |
| 3 | 5 | paraffin oil | 4 | 1 | -3 | |
| 3 | 5 | ethyl (2*E*,4*Z*)-decadienoate | 1 | 4 | 3 | 6 |
| 3 | 5 | *p*-cresol | 0 | 0 | 0 | 3 |
| 3 | 5 | *β*-bisabolene | 0 | 15 | 15 | 18 |
| 3 | 5 | (*E*)-myrcenol | 5 | 5 | 0 | 3 |
| 3 | 5 | nonan-1-ol | 0 | 1 | 1 | 4 |
| 3 | 5 | isoamyl acetate | 4 | 1 | -3 | 0 |
| 3 | 5 | 2-phenylethyl acetate | 3 | 3 | 0 | 3 |
| 3 | 5 | octan-1-ol | 9 | 6 | -3 | 0 |
| 3 | 5 | benzyl alcohol | 5 | 0 | -5 | -2 |
| 3 | 5 | methyl jasmonate | 0 | 1 | 1 | 4 |
| 3 | 5 | (*E*)-*β*-ocimene | 2 | 0 | -2 | 1 |
| 3 | 5 | eucarvone | 9 | 7 | -2 | 1 |
| 3 | 5 | *α*-camphorene | 11 | 4 | -7 | -4 |
| 3 | 5 | (-)-trans-pinocarveol | 0 | 0 | 0 | 3 |
| 3 | 5 | cryptone | 3 | 2 | -1 | 2 |
| 3 | 5 | (+)-cis-carveol | 4 | 1 | -3 | 0 |

**Table S9.** SSR responses to Panel 4 for PsimOR14. Related to Fig. 3.

| Panel | Replicate | Compound | Pre-stimulation spikes | Post-stimulation spikes | Generated spikes | Δ Spikes/s |
|---|---|---|---|---|---|---|
| 4 | 1 | paraffin oil | 6 | 1 | -5 | |
| 4 | 1 | (-)-verbenone | 14 | 4 | -10 | -5 |
| 4 | 1 | oct-1-en-3-ol | 4 | 2 | -2 | 3 |
| 4 | 1 | acetophenone | 12 | 20 | 8 | 13 |
| 4 | 1 | 4-vinylanisole | 12 | 14 | 2 | 7 |
| 4 | 1 | 4-ethylguaiacol | 6 | 2 | -4 | 1 |
| 4 | 1 | (±)-camphor | 10 | 6 | -4 | 1 |
| 4 | 1 | octan-1-ol | 16 | 6 | -10 | -5 |
| 4 | 1 | methyleugenol | 18 | 2 | -16 | -11 |
| 4 | 1 | styrene | 16 | 14 | -2 | 3 |
| 4 | 1 | benzaldehyde | 20 | 6 | -14 | -9 |
| 4 | 1 | linalool | 21 | 0 | -21 | -16 |
| 4 | 1 | 4-methylanisole | 20 | 6 | -14 | -9 |
| 4 | 1 | 2-methylbutan-1-ol | 4 | 0 | -4 | 1 |
| 4 | 1 | isoamyl alcohol | 12 | 0 | -12 | -7 |
| 4 | 1 | 2-methylbutyl acetate | 18 | 14 | -4 | 1 |
| 4 | 1 | (±)-myrtenol | 7 | 2 | -5 | 0 |
| 4 | 2 | paraffin oil | 7 | 4 | -3 | |
| 4 | 2 | (-)-verbenone | 2 | 12 | 10 | 13 |
| 4 | 2 | oct-1-en-3-ol | 7 | 8 | 1 | 4 |
| 4 | 2 | acetophenone | 4 | 10 | 6 | 9 |
| 4 | 2 | 4-vinylanisole | 8 | 5 | -3 | 0 |
| 4 | 2 | 4-ethylguaiacol | 6 | 10 | 4 | 7 |
| 4 | 2 | (±)-camphor | 4 | 11 | 7 | 10 |
| 4 | 2 | octan-1-ol | 13 | 8 | -5 | -2 |
| 4 | 2 | methyleugenol | 5 | 14 | 9 | 12 |
| 4 | 2 | styrene | 10 | 6 | -4 | -1 |
| 4 | 2 | benzaldehyde | 7 | 11 | 4 | 7 |
| 4 | 2 | linalool | 5 | 5 | 0 | 3 |
| 4 | 2 | 4-methylanisole | 10 | 4 | -6 | -3 |
| 4 | 2 | 2-methylbutan-1-ol | 8 | 6 | -2 | 1 |
| 4 | 2 | isoamyl alcohol | 11 | 12 | 1 | 4 |
| 4 | 2 | 2-methylbutyl acetate | 10 | 10 | 0 | 3 |
| 4 | 2 | (±)-myrtenol | 7 | 9 | 2 | 5 |
| 4 | 2 | 2-phenylethanol | 4 | 8 | 4 | 7 |
| 4 | 2 | hexan-1-ol | 9 | 4 | -5 | -2 |
| 4 | 2 | 2,3-dihydrobenzofuran | 9 | 4 | -5 | -2 |
| 4 | 2 | geranyl acetone | 8 | 6 | -2 | 1 |
| 4 | 3 | paraffin oil | 4 | 2 | -2 | |
| 4 | 3 | (-)-verbenone | 12 | 8 | -4 | -2 |
| 4 | 3 | oct-1-en-3-ol | 7 | 5 | -2 | 0 |
| 4 | 3 | acetophenone | 2 | 10 | 8 | 10 |
| 4 | 3 | 4-vinylanisole | 8 | 12 | 4 | 6 |
| 4 | 3 | 4-ethylguaiacol | 1 | 11 | 10 | 12 |
| 4 | 3 | (±)-camphor | 1 | 10 | 9 | 11 |
| 4 | 3 | octan-1-ol | 9 | 6 | -3 | -1 |
| 4 | 3 | methyleugenol | 9 | 6 | -3 | -1 |
| 4 | 3 | styrene | 3 | 5 | 2 | 4 |
| 4 | 3 | benzaldehyde | 11 | 6 | -5 | -3 |
| 4 | 3 | linalool | 10 | 1 | -9 | -7 |
| 4 | 3 | 4-methylanisole | 3 | 9 | 6 | 8 |
| 4 | 3 | 2-methylbutan-1-ol | 7 | 2 | -5 | -3 |
| 4 | 3 | isoamyl alcohol | 10 | 9 | -1 | 1 |
| 4 | 3 | 2-methylbutyl acetate | 10 | 6 | -4 | -2 |
| 4 | 3 | (±)-myrtenol | 5 | 3 | -2 | 0 |
| 4 | 3 | 2-phenylethanol | 2 | 8 | 6 | 8 |
| 4 | 3 | hexan-1-ol | 7 | 4 | -3 | -1 |
| 4 | 3 | 2,3-dihydrobenzofuran | 6 | 9 | 3 | 5 |
| 4 | 3 | geranyl acetone | 6 | 9 | 3 | 5 |
| 4 | 4 | paraffin oil | 10 | 6 | -4 | |
| 4 | 4 | (-)-verbenone | 11 | 5 | -6 | -2 |
| 4 | 4 | oct-1-en-3-ol | 7 | 8 | 1 | 5 |
| 4 | 4 | acetophenone | 13 | 3 | -10 | -6 |
| 4 | 4 | 4-vinylanisole | 8 | 8 | 0 | 4 |
| 4 | 4 | 4-ethylguaiacol | 7 | 6 | -1 | 3 |
| 4 | 4 | (±)-camphor | 1 | 1 | 0 | 4 |
| 4 | 4 | octan-1-ol | 7 | 7 | 0 | 4 |
| 4 | 4 | methyleugenol | 14 | 7 | -7 | -3 |
| 4 | 4 | styrene | 11 | 6 | -5 | -1 |
| 4 | 4 | benzaldehyde | 5 | 14 | 9 | 13 |
| 4 | 4 | linalool | 8 | 0 | -8 | -4 |
| 4 | 4 | 4-methylanisole | 7 | 5 | -2 | 2 |
| 4 | 4 | 2-methylbutan-1-ol | 5 | 0 | -5 | -1 |
| 4 | 4 | isoamyl alcohol | 9 | 3 | -6 | -2 |
| 4 | 4 | 2-methylbutyl acetate | 6 | 9 | 3 | 7 |
| 4 | 4 | (±)-myrtenol | 3 | 6 | 3 | 7 |
| 4 | 4 | 2-phenylethanol | 6 | 5 | -1 | 3 |
| 4 | 4 | hexan-1-ol | 9 | 8 | -1 | 3 |
| 4 | 4 | 2,3-dihydrobenzofuran | 6 | 4 | -2 | 2 |
| 4 | 4 | geranyl acetone | 10 | 7 | -3 | 1 |
| 4 | 5 | paraffin oil | 8 | 7 | -1 | |
| 4 | 5 | (-)-verbenone | 6 | 3 | -3 | -2 |
| 4 | 5 | oct-1-en-3-ol | 3 | 1 | -2 | -1 |
| 4 | 5 | acetophenone | 6 | 2 | -4 | -3 |
| 4 | 5 | 4-vinylanisole | 5 | 3 | -2 | -1 |
| 4 | 5 | 4-ethylguaiacol | 9 | 5 | -4 | -3 |
| 4 | 5 | (±)-camphor | 8 | 5 | -3 | -2 |
| 4 | 5 | octan-1-ol | 6 | 4 | -2 | -1 |
| 4 | 5 | methyleugenol | 7 | 8 | 1 | 2 |
| 4 | 5 | styrene | 7 | 6 | -1 | 0 |
| 4 | 5 | benzaldehyde | 10 | 5 | -5 | -4 |
| 4 | 5 | linalool | 9 | 2 | -7 | -6 |
| 4 | 5 | 4-methylanisole | 10 | 9 | -1 | 0 |
| 4 | 5 | 2-methylbutan-1-ol | 10 | 4 | -6 | -5 |
| 4 | 5 | isoamyl alcohol | 10 | 7 | -3 | -2 |
| 4 | 5 | 2-methylbutyl acetate | 10 | 8 | -2 | -1 |
| 4 | 5 | (±)-myrtenol | 6 | 5 | -1 | 0 |
| 4 | 5 | 2-phenylethanol | 8 | 7 | -1 | 0 |
| 4 | 5 | hexan-1-ol | 9 | 5 | -4 | -3 |
| 4 | 5 | 2,3-dihydrobenzofuran | 7 | 6 | -1 | 0 |
| 4 | 5 | geranyl acetone | 9 | 6 | -3 | -2 |
| 4 | 6 | paraffin oil | 9 | 10 | 1 | |
| 4 | 6 | (-)-verbenone | 7 | 4 | -3 | -4 |
| 4 | 6 | oct-1-en-3-ol | 6 | 8 | 2 | 1 |
| 4 | 6 | acetophenone | 12 | 6 | -6 | -7 |
| 4 | 6 | 4-vinylanisole | 7 | 6 | -1 | -2 |
| 4 | 6 | 4-ethylguaiacol | 8 | 5 | -3 | -4 |
| 4 | 6 | (±)-camphor | 11 | 14 | 3 | 2 |
| 4 | 6 | octan-1-ol | 13 | 5 | -8 | -9 |
| 4 | 6 | methyleugenol | 5 | 6 | 1 | 0 |
| 4 | 6 | styrene | 9 | 6 | -3 | -4 |
| 4 | 6 | benzaldehyde | 6 | 7 | 1 | 0 |
| 4 | 6 | linalool | 11 | 1 | -10 | -11 |
| 4 | 6 | 4-methylanisole | 9 | 8 | -1 | -2 |
| 4 | 6 | 2-methylbutan-1-ol | 10 | 4 | -6 | -7 |
| 4 | 6 | isoamyl alcohol | 7 | 7 | 0 | -1 |
| 4 | 6 | 2-methylbutyl acetate | 6 | 7 | 1 | 0 |
| 4 | 6 | (±)-myrtenol | 7 | 6 | -1 | -2 |
| 4 | 6 | 2-phenylethanol | 10 | 8 | -2 | -3 |
| 4 | 6 | hexan-1-ol | 11 | 6 | -5 | -6 |
| 4 | 6 | 2,3-dihydrobenzofuran | 12 | 9 | -3 | -4 |
| 4 | 6 | geranyl acetone | 12 | 9 | -3 | -4 |

**Table S10.** SSR responses to Panel 1 by neocembrene sensillum in *P. simplex* workers. Related to Fig. 5.

| Replicate | Compound | Pre-stimulation spikes | Post-stimulation spikes | Generated spikes | Δ Spikes/s |
|---|---|---|---|---|---|
| 1 | hexane | 8 | 8 | 0 | |
| 1 | (3*Z*,6*Z*)-dodecadien-1-ol | 5 | 8 | 3 | 6 |
| 1 | (3*Z*)-dodecen-1-ol | 12 | 21 | 9 | 18 |
| 1 | (3*Z*,6*Z*,8*E*)-dodecatrien-1-ol | 10 | 13 | 3 | 6 |
| 1 | *n*-eicosane | 6 | 8 | 2 | 4 |
| 1 | 1-octadecanol | 5 | 8 | 3 | 6 |
| 1 | *n*-docosane | 3 | 5 | 2 | 4 |
| 1 | (3*R*,6*E*)-nerolidol | 4 | 16 | 12 | 24 |
| 1 | neocembrene | 11 | 34 | 23 | 46 |
| 1 | δ-cadinene | 9 | 12 | 3 | 6 |
| 1 | dodec-3-yn-1-ol | 5 | 6 | 1 | 2 |
| 1 | geranylgeraniol | 4 | 12 | 8 | 16 |
| 2 | hexane | 18 | 18 | 0 | |
| 2 | (3*Z*,6*Z*)-dodecadien-1-ol | 9 | 11 | 2 | 4 |
| 2 | (3*Z*)-dodecen-1-ol | 2 | 5 | 3 | 6 |
| 2 | (3*Z*,6*Z*,8*E*)-dodecatrien-1-ol | 8 | 6 | -2 | -4 |
| 2 | neocembrene | 3 | 41 | 38 | 76 |
| 2 | geranylgeraniol | 2 | 11 | 9 | 18 |
| 3 | hexane | 6 | 6 | 0 | |
| 3 | (3*Z*,6*Z*)-dodecadien-1-ol | 10 | 12 | 2 | 4 |
| 3 | (3*Z*)-dodecen-1-ol | 15 | 9 | -6 | -12 |
| 3 | (3*Z*,6*Z*,8*E*)-dodecatrien-1-ol | 9 | 11 | 2 | 4 |
| 3 | neocembrene | 10 | 46 | 36 | 72 |
| 3 | geranylgeraniol | 6 | 16 | 10 | 20 |
| 3 | δ-cadinene | 12 | 13 | 1 | 2 |
| 4 | hexane | 5 | 6 | 1 | |
| 4 | (3*Z*,6*Z*)-dodecadien-1-ol | 10 | 12 | 2 | 2 |
| 4 | (3*Z*)-dodecen-1-ol | 10 | 2 | -8 | -18 |
| 4 | (3*Z*,6*Z*,8*E*)-dodecatrien-1-ol | 9 | 9 | 0 | -2 |
| 4 | neocembrene | 7 | 45 | 38 | 74 |
| 4 | geranylgeraniol | 3 | 9 | 6 | 10 |
| 5 | hexane | 6 | 10 | 4 | |
| 5 | (3*Z*,6*Z*)-dodecadien-1-ol | 7 | 13 | 6 | 4 |
| 5 | (3*Z*)-dodecen-1-ol | 6 | 15 | 9 | 10 |
| 5 | (3*Z*,6*Z*,8*E*)-dodecatrien-1-ol | 9 | 11 | 2 | -4 |
| 5 | neocembrene | 5 | 51 | 46 | 84 |
| 6 | hexane | 12 | 4 | -8 | |
| 6 | (3*Z*,6*Z*)-dodecadien-1-ol | 7 | 14 | 7 | 30 |
| 6 | (3*Z*)-dodecen-1-ol | 11 | 7 | -4 | 8 |
| 6 | (3*Z*,6*Z*,8*E*)-dodecatrien-1-ol | 10 | 18 | 8 | 32 |
| 6 | neocembrene | 3 | 48 | 45 | 106 |
| 6 | geranylgeraniol | 9 | 14 | 5 | 26 |
| 7 | hexane | 36 | 33 | -3 | |
| 7 | (3*Z*,6*Z*)-dodecadien-1-ol | 40 | 53 | 13 | 32 |
| 7 | (3*Z*)-dodecen-1-ol | 47 | 49 | 2 | 10 |
| 7 | (3*Z*,6*Z*,8*E*)-dodecatrien-1-ol | 55 | 64 | 9 | 24 |
| 7 | *n*-eicosane | 54 | 54 | 0 | 6 |
| 7 | 1-octadecanol | 54 | 49 | -5 | -4 |
| 7 | *n*-docosane | 56 | 49 | -7 | -8 |
| 7 | (3*R*,6*E*)-nerolidol | 57 | 50 | -7 | -8 |
| 7 | δ-cadinene | 52 | 46 | -6 | -6 |
| 7 | dodec-3-yn-1-ol | 48 | 54 | 6 | 18 |
| 7 | neocembrene | 40 | 74 | 34 | 74 |
| 7 | geranylgeraniol | 17 | 40 | 23 | 52 |
| 8 | hexane | 34 | 32 | -2 | |
| 8 | neocembrene | 18 | 46 | 28 | 60 |
| 8 | geranylgeraniol | 20 | 39 | 19 | 42 |
| 9 | hexane | 65 | 70 | 5 | |
| 9 | (3*Z*,6*Z*)-dodecadien-1-ol | 28 | 33 | 5 | 0 |
| 9 | (3*Z*)-dodecen-1-ol | 37 | 36 | -1 | -12 |
| 9 | (3*Z*,6*Z*,8*E*)-dodecatrien-1-ol | 29 | 27 | -2 | -14 |
| 9 | *n*-eicosane | 36 | 42 | 6 | 2 |
| 9 | 1-octadecanol | 35 | 34 | -1 | -12 |
| 9 | *n*-docosane | 34 | 35 | 1 | -8 |
| 9 | (3*R*,6*E*)-nerolidol | 19 | 30 | 11 | 12 |
| 9 | neocembrene | 22 | 54 | 32 | 54 |
| 9 | δ-cadinene | 19 | 15 | -4 | -18 |
| 9 | dodec-3-yn-1-ol | 35 | 31 | -4 | -18 |
| 9 | geranylgeraniol | 20 | 44 | 24 | 38 |
| 10 | hexane | 17 | 22 | 5 | |
| 10 | neocembrene | 20 | 39 | 19 | 28 |
| 10 | geranylgeraniol | 17 | 41 | 24 | 38 |
| 11 | hexane | 6 | 11 | 5 | |
| 11 | neocembrene | 19 | 46 | 27 | 44 |
| 11 | geranylgeraniol | 24 | 34 | 10 | 10 |
| 12 | hexane | 28 | 26 | -2 | |
| 12 | neocembrene | 31 | 56 | 25 | 54 |
| 13 | hexane | 41 | 37 | -4 | |
| 13 | neocembrene | 17 | 38 | 21 | 50 |
| 13 | geranylgeraniol | 8 | 28 | 20 | 48 |
| 13 | (3*Z*,6*Z*)-dodecadien-1-ol | 15 | 13 | -2 | 4 |
| 13 | (3*Z*)-dodecen-1-ol | 18 | 20 | 2 | 12 |
| 13 | (3*Z*,6*Z*,8*E*)-dodecatrien-1-ol | 12 | 13 | 1 | 10 |
| 13 | *n*-eicosane | 15 | 18 | 3 | 14 |
| 13 | 1-octadecanol | 7 | 11 | 4 | 16 |
| 13 | *n*-docosane | 7 | 11 | 4 | 16 |
| 13 | (3*R*,6*E*)-nerolidol | 27 | 19 | -8 | -8 |
| 13 | δ-cadinene | 27 | 23 | -4 | 0 |
| 13 | dodec-3-yn-1-ol | 18 | 30 | 12 | 32 |
| 14 | hexane | 25 | 21 | -4 | |
| 14 | neocembrene | 18 | 68 | 50 | 108 |
| 14 | geranylgeraniol | 15 | 22 | 7 | 22 |
| 14 | (3*Z*,6*Z*)-dodecadien-1-ol | 28 | 15 | -13 | -18 |
| 14 | (3*Z*)-dodecen-1-ol | 32 | 26 | -6 | -4 |
| 14 | (3*Z*,6*Z*,8*E*)-dodecatrien-1-ol | 20 | 24 | 4 | 16 |
| 14 | *n*-eicosane | 21 | 17 | -4 | 0 |
| 14 | 1-octadecanol | 30 | 23 | -7 | -6 |
| 14 | *n*-docosane | 27 | 24 | -3 | 2 |
| 14 | (3*R*,6*E*)-nerolidol | 7 | 9 | 2 | 12 |
| 14 | δ-cadinene | 13 | 11 | -2 | 4 |
| 14 | dodec-3-yn-1-ol | 8 | 8 | 0 | 8 |
| 15 | hexane | 9 | 8 | -1 | |
| 15 | (3*Z*,6*Z*)-dodecadien-1-ol | 24 | 21 | -3 | -4 |
| 15 | (3*Z*)-dodecen-1-ol | 24 | 29 | 5 | 12 |
| 15 | (3*Z*,6*Z*,8*E*)-dodecatrien-1-ol | 13 | 14 | 1 | 4 |
| 15 | *n*-eicosane | 8 | 19 | 11 | 24 |
| 15 | 1-octadecanol | 23 | 19 | -4 | -6 |
| 15 | *n*-docosane | 30 | 26 | -4 | -6 |
| 15 | (3*R*,6*E*)-nerolidol | 9 | 16 | 7 | 16 |
| 15 | neocembrene | 12 | 40 | 28 | 58 |
| 15 | δ-cadinene | 9 | 9 | 0 | 2 |
| 15 | dodec-3-yn-1-ol | 16 | 8 | -8 | -14 |
| 15 | geranylgeraniol | 10 | 24 | 14 | 30 |
| 16 | hexane | 13 | 7 | -6 | |
| 16 | neocembrene | 18 | 38 | 20 | 52 |
| 16 | geranylgeraniol | 9 | 26 | 17 | 46 |
| 16 | (3*Z*,6*Z*)-dodecadien-1-ol | 15 | 12 | -3 | 6 |
| 16 | (3*Z*)-dodecen-1-ol | 12 | 25 | 13 | 38 |
| 16 | (3*Z*,6*Z*,8*E*)-dodecatrien-1-ol | 11 | 13 | 2 | 16 |
| 16 | *n*-eicosane | 15 | 18 | 3 | 18 |
| 16 | 1-octadecanol | 7 | 13 | 6 | 24 |
| 16 | *n*-docosane | 9 | 10 | 1 | 14 |
| 16 | (3*R*,6*E*)-nerolidol | 7 | 13 | 6 | 24 |
| 16 | δ-cadinene | 12 | 13 | 1 | 14 |
| 16 | dodec-3-yn-1-ol | 8 | 18 | 10 | 32 |
| 17 | hexane | 16 | 14 | -2 | |
| 17 | neocembrene | 7 | 32 | 25 | 54 |
| 17 | geranylgeraniol | 8 | 14 | 6 | 16 |
| 17 | (3*Z*,6*Z*)-dodecadien-1-ol | 10 | 7 | -3 | -2 |
| 17 | (3*Z*)-dodecen-1-ol | 18 | 8 | -10 | -16 |
| 17 | (3*Z*,6*Z*,8*E*)-dodecatrien-1-ol | 17 | 9 | -8 | -12 |
| 17 | *n*-eicosane | 7 | 8 | 1 | 6 |
| 17 | 1-octadecanol | 17 | 9 | -8 | -12 |
| 17 | *n*-docosane | 14 | 10 | -4 | -4 |
| 17 | (3*R*,6*E*)-nerolidol | 8 | 8 | 0 | 4 |
| 17 | δ-cadinene | 9 | 14 | 5 | 14 |
| 17 | dodec-3-yn-1-ol | 9 | 10 | 1 | 6 |

**Table S11.** SSR dose response data for neocembrene and *P. simplex* neocembrene sensillum. Related to Fig. 5.

| | | Δ Spikes/s | | | | | | | | | | |
|---|---|---|---|---|---|---|---|---|---|---|---|---|
| | replicate | 1 | 2 | 3 | 4 | 5 | 6 | 7 | 8 | 9 | 10 | 11 |
| Dose | 0.01 ng | 62 | 44 | | 18 | | 14 | 8 | 30 | 10 | 4 | 18 |
| | 0.1 ng | 76 | 24 | 20 | 64 | 90 | 22 | 42 | 36 | 12 | 10 | 42 |
| | 1 ng | 90 | 46 | 54 | 86 | 106 | 14 | 30 | 38 | 20 | 12 | 36 |
| | 10 ng | 84 | 44 | 50 | 88 | 120 | 22 | 14 | | 16 | 28 | 46 |
| | 100 ng | 104 | 54 | 34 | 112 | 124 | 36 | 8 | | 36 | 38 | 40 |

**Table S12.** Nucleotide and protein sequences of PsimOR14. Related to Fig. 6.

***PsimOR14***

```
ATGATTCGATCAAAGAGAAAGGAGAGCCAAGCAAACGAAACAAAACACACATTAACAAGTGAAGAGCAACCTTCTGATTG
TGACGTGAAGATCATGACACTCAGTATTATGCTGAATGCGGCTGGCCTCCTACCTCCAGCCAAATCGTCATCATTGATCA
GACTGGCCTACAAAGTATTTGTAGTATTTATTCACATACTTTTCGTCTTAACGCTGATAGGACAGATAATGGCAGTAGTG
GTTTACTGGGGAGACATTCCTCTAATTGCAACCACAATAAGCTTGATGACTAGTCTGATTGGATCGATGAGTTCATCCAT
AAATTTTCTTCTAAACAGAAAGAAGTACATGCGTCTTGCGGACACGTTGAAAACAGAATTTGTTGCCAAATTGAAATCAA
AATATATCAAAATTATTTTAAATGCTGAACGTCAGGTTGTATTCTGTGGGATACTCGTATGTATTGTAGCTGTATGTATT
GGATTTATTTGGATAGTCGTGCCATTTTTAAGTACCAACACCCCATTTGACTTTGCAAATGAAAAAAGTGTCAAAGAAGG
AAGCCGCATGGAGGAATTAATTCTTGTGATGTGGCTCCCTTCTAAATTTGAACAGTCCCCTCAATTTGAAATAATAGTTT
TTTTACAAATCTTTGTCGTAACGTTTGCATTAGCAATGATCTATTCAGTTGATATGATGTTACTATCTCTGATGAGCCAC
GCTGCTGCACAGTTCAGGGTTTTGAATGCCATGCTGAATGACATGCACGAAAATGTTCGTGAAGACGAGATTCACAGAAC
AAGAAACATGGCTTCATTGGTCACTGGCACTGACATCTCGTACATGGAGTTCTCTTCTACCAATTCTTGGAATGGAAACA
CAGAGCATTCTGGAAGCGCTGGTGTTGAGTTGGACAGCCTAAAAAATGAAGACTGTGAAGAAGATCCTGTCCGACAGTAC
CTCGTTGAGTGCATTAGATATCACCAGGCTGTAATTGAGTTTGTTGACCAACTGAACGAGGTGTTCGGCGCAGTGAGCTT
CGTGAAGATGCTTGACTGGCCTTTTGCGATTTGTATGACAGGATTTCAGTTGACACAGACTGTAGAGAGCCAGGAGGATT
TACTTAAATTCATCTCCCTGTTTGCTGGGGTTGTATATCTAATAATCTCTTACATTTGGTTCGGACAGCAAGTAATTGAC
GAGAGCGAGGAAGTAGCAACAGCGTTGTACAGCACTGACTGGTACAACCAGGCACCAGGGTTCAAACGTCTGCTGCCTGT
AGCCATCATGCGGGCTTCGAATTCTGTCAAAGTTAAAGCTGGGGTGTTCTTTGACATGTCTTGCGTCACGTTAGCTTCGA
TTATGAATGCATCTTACACGTATTTTATGATGCTAATTCATCTACACGACTCCTAA
```

**PsimOR14**

```
MIRSKRKESQANETKHTLTSEEQPSDCDVKIMTLSIMLNAAGLLPPAKSSSLIRLAYKVFVVFIHILFVLTLIGQIMAVV
VYWGDIPLIATTISLMTSLIGSMSSSINFLLNRKKYMRLADTLKTEFVAKLKSKYIKIILNAERQVVFCGILVCIVAVCI
GFIWIVVPFLSTNTPFDFANEKSVKEGSRMEELILVMWLPSKFEQSPQFEIIVFLQIFVVTFALAMIYSVDMMLLSLMSH
AAAQFRVLNAMLNDMHENVREDEIHRTRNMASLVTGTDISYMEFSSTNSWNGNTEHSGSAGVELDSLKNEDCEEDPVRQY
LVECIRYHQAVIEFVDQLNEVFGAVSFVKMLDWPFAICMTGFQLTQTVESQEDLLKFISLFAGVVYLIISYIWFGQQVID
ESEEVATALYSTDWYNQAPGFKRLLPVAIMRASNSVKVKAGVFFDMSCVTLASIMNASYTYFMMLIHLHDSX
```

**Table S13.** MM/PBSA calculated interaction energies of ligands with PsimOR14 decomposed into per-residue contributions.

| Interaction energy (kcal/mol) | neocembrene | geranylgeraniol | (+)-limonene |
|---|---|---|---|
| | | mean ± SD | |
| Cys154 | -0.90 ± 0.58 | -0.71 ± 0.35 | -0.17 ± 0.27 |
| Ala157 | -0.68 ± 0.46 | -1.02 ± 0.38 | -0.45 ± 0.38 |
| Val158 | -1.16 ± 0.77 | -1.72 ± 0.56 | -0.23 ± 0.44 |
| Gly161 | × | -0.82 ± 0.38 | × |
| Phe162 | × | -1.08 ± 0.67 | × |
| Ile165 | × | -0.66 ± 0.49 | × |
| Ile217 | × | -1.03 ± 0.47 | × |
| Val220 | × | -0.72 ± 0.31 | × |
| Thr221 | -1.00 ± 0.53 | -1.19 ± 0.47 | -0.17 ± 0.35 |
| Leu224 | -1.09 ± 0.58 | -1.41 ± 0.39 | -0.78 ± 0.46 |
| Ala225 | -0.93 ± 0.6 | -0.97 ± 0.45 | -0.17 ± 0.29 |
| Tyr228 | -1.10 ± 0.72 | -1.06 ± 0.49 | -0.47 ± 0.4 |

**Table S14.** Differential expression of *P. simplex* ORs (DESeq2 and edge R analyses), based on heads (including antennae) of workers and soldiers (three replicates). Related to Fig. 7.

| | DESeq2 | | | edgeR | |
|---|---|---|---|---|---|
| OR | $log_2$ fold change (worker vs soldier) | lfcSE | p-value | $log_2$ fold change (worker vs soldier) | p-value |
| PsimOR1 | -0.5905 | 0.3426 | 0.0848 | -0.6679 | 0.0186 |
| PsimOR10 | -0.4006 | 0.9211 | 0.6636 | -0.4375 | 0.5905 |
| PsimOR11 | 0.3565 | 0.5005 | 0.4763 | 0.262 | 0.5589 |
| PsimOR12 | -0.2703 | 1.2658 | 0.8309 | -0.4127 | 0.8431 |
| PsimOR13 | 0.1677 | 0.6478 | 0.7957 | 0.0539 | 1 |
| PsimOR14 | -1.8565 | 0.8841 | 0.0357 | -1.8989 | 0.0036 |
| PsimOR15 | -0.578 | 0.8735 | 0.5082 | -0.6829 | 0.4084 |
| PsimOR16 | -0.4973 | 1.1753 | 0.6722 | -0.5878 | 0.613 |
| PsimOR17 | -0.7328 | 3.7695 | 0.8459 | -0.5892 | 1 |
| PsimOR18 | 1.2163 | 1.6983 | 0.4739 | 1.0486 | 0.5486 |
| PsimOR19 | 0.6517 | 1.0553 | 0.5369 | 0.5408 | 0.5589 |
| PsimOR2 | 1.2943 | 0.7769 | 0.0957 | 1.1238 | 0.1342 |
| PsimOR20 | -2.9803 | 0.7045 | 0 | -3.0603 | 0 |
| PsimOR22 | 0.5913 | 1.1359 | 0.6027 | 0.5019 | 0.7071 |
| PsimOR23 | -0.0816 | 1.3315 | 0.9511 | -0.1249 | 1 |
| PsimOR24 | -1.3934 | 0.8105 | 0.0856 | -1.4649 | 0.025 |
| PsimOR25 | 0.9132 | 1.4392 | 0.5257 | 0.6897 | 0.6457 |
| PsimOR26 | -0.879 | 0.1926 | 0 | -0.9787 | 0 |
| PsimOR27 | 0.2988 | 1.1656 | 0.7977 | 0.2 | 1 |
| PsimOR28 | 0.3981 | 1.1226 | 0.7229 | 0.3443 | 0.722 |
| PsimOR29 | -0.4531 | 1.3799 | 0.7426 | -0.45 | 0.6812 |
| PsimOR3 | -0.037 | 1.156 | 0.9744 | -0.1663 | 0.8475 |
| PsimOR30 | 0.7053 | 1.184 | 0.5514 | 0.5977 | 0.6758 |
| PsimOR31 | 0.1924 | 0.5699 | 0.7357 | 0.0691 | 0.9348 |
| PsimOR32 | -0.2186 | 1.4043 | 0.8763 | -0.288 | 1 |
| PsimOR34 | -0.4167 | 0.4244 | 0.3261 | -0.5383 | 0.1738 |
| PsimOR35 | -1.0587 | 2.6395 | 0.6883 | -0.8036 | 0.5553 |
| PsimOR36 | -0.3664 | 0.9102 | 0.6873 | -0.5093 | 0.5249 |
| PsimOR37 | 0.3495 | 1.2265 | 0.7756 | 0.2614 | 0.8477 |
| PsimOR39 | -0.0669 | 0.6154 | 0.9135 | -0.1886 | 0.7502 |
| PsimOR4 | 0.0586 | 0.7043 | 0.9336 | -0.0669 | 1 |
| PsimOR40 | -0.5383 | 0.34 | 0.1134 | -0.6076 | 0.0395 |
| PsimOR41 | -0.3886 | 0.2793 | 0.164 | -0.5129 | 0.0916 |
| PsimOR42 | NA | NA | NA | 0 | 1 |
| PsimOR43 | -0.5866 | 1.4600 | 0.6878 | -0.5596 | 0.7796 |
| PsimOR44 | 0.1981 | 1.6775 | 0.906 | 0.128 | 1 |
| PsimOR45 | -1.5455 | 1.331 | 0.2456 | -1.4406 | 0.1817 |
| PsimOR46 | -3.0579 | 1.863 | 0.1007 | -2.9 | 0.0445 |
| PsimOR47 | 2.375 | 1.7303 | 0.1699 | 2.2464 | 0.1134 |
| PsimOR48 | 0.0016 | 1.2033 | 0.9989 | -0.0757 | 1 |
| PsimOR49 | -1.0692 | 1.6425 | 0.5151 | -0.9962 | 0.3814 |
| PsimOR5 | 0.4942 | 0.6163 | 0.4227 | 0.4152 | 0.4609 |
| PsimOR50 | 0.229 | 4.0805 | 0.9553 | 1.6045 | 1 |
| PsimOR6 | -0.0086 | 0.5823 | 0.9882 | -0.0515 | 1 |
| PsimOR7 | 0.161 | 0.4781 | 0.7363 | 0.1071 | 0.8517 |
| PsimOR8 | 0.4963 | 0.6856 | 0.4692 | 0.3467 | 0.6466 |
| PsimOR9 | 2.0761 | 0.9969 | 0.0373 | 1.8932 | 0.0303 |

**Table S15.** Differential sensitivity of *P. simplex* workers and soldiers to neocembrene inferred from EAG responses to the dose of 10 ng. Related to Fig. 7.

| EAG response normalized to air stimulation | | $log_2$ value | |
|---|---|---|---|
| worker | soldier | worker | soldier |
| 199.52 | 92.67 | 7.64 | 6.53 |
| 322.81 | 116.73 | 8.33 | 6.87 |
| 132.44 | 136.93 | 7.05 | 7.10 |
| 142.77 | 78.81 | 7.16 | 6.30 |
| 124.07 | 120.01 | 6.96 | 6.91 |
| 135.73 | 120.49 | 7.08 | 6.91 |
| 192.41 | 136.59 | 7.59 | 7.09 |
| 146.41 | 116.43 | 7.19 | 6.86 |
| 122.17 | 176.51 | 6.93 | 7.46 |
| 201.83 | 122.48 | 7.66 | 6.94 |
| 123.71 | 110.96 | 6.95 | 6.79 |
| 142.15 | 139.64 | 7.15 | 7.13 |
| 115.99 | 138.32 | 6.86 | 7.11 |
| 164.7 | 132.45 | 7.36 | 7.05 |
| 167.08 | 152.31 | 7.38 | 7.25 |

**Table S16.** List of primers.

| Primer Name | Sequence (5' to 3' direction) | Use |
|---|---|---|
| PsimOR14_F | ATGATTCGATCAAAGAGAAAGG | Cloning |
| PsimOR14_R | TTAGGAGTCGTGTAGATGAAT | Cloning |
| PsimO31_F | ATGGAATACATAAAAAATGAAACATATTCTCA | Cloning |
| PsimO31_R | TCAACCTACGACATGTGAGTTATT | Cloning |
| PsimOR9_F | ATGGACAGCCTTTACGACCAATCTT | Cloning |
| PsimOR9_R | TCATTCAGTGACTGAGGGATCCTT | Cloning |
| PsimO30_F | ATGGAGCACAGGAAATACAAAGTGACAA | Cloning |
| PsimO30_R | TTACGTTCCCTGATTTGTGTCGGTAT | Cloning |
| PsimOrco_F | ATGTACAAGTTCAGGTTACACG | cDNA check |
| PsimOrco_R | CTAGTTGAGCTGTACCAACAC | cDNA check |
| GW1 | GTTGCAACAAATTGATGAGCAATGC | Sanger Sequencing & Colony PCR |
| GW2 | GTTGCAACAAATTGATGAGCAATTA | Sanger Sequencing & Colony PCR |
| UAS1 | TAGCGAGCGCCGGAGTATAAATAG | Sanger Sequencing |
| UAS2 | ACTGATTTCGACGGTTACCC | Sanger Sequencing |
| DmOr22a_F | TCTCCAGCATCGCCGAGTGT | Single-Wing PCR |
| DmOr22a_R | CGGCAGAGGTCCAGTCCGAT | Single-Wing PCR |
| PsimOR14_SW_F | GAGAGCCAAGCAAACGAAAC | Single-Wing PCR |
| PsimOR14_SW_R | TTTAGAAGGGAGCCACATCAC | Single-Wing PCR |
| PsimO31_SW_F | GCTGGGTTAATCCCGATCAT | Single-Wing PCR |
| PsimO31_SW_R | GCATGGCACCAAATAGTTCTTC | Single-Wing PCR |
| PsimOR9_SW_F | TGGGCGAAACTGAGGATATG | Single-Wing PCR |
| PsimOR9_SW_R | CGAGCCGACATAGAAGAAGAG | Single-Wing PCR |
| PsimO30_SW_F | TGCCATCACCAGCAGATAAA | Single-Wing PCR |
| PsimO30_SW_R | CACCGACTGACTCAGCATATT | Single-Wing PCR |

**Table S17.** Characteristics and origin of *D. melanogaster* lines used.

| Fly line | Source |
|---|---|
| **$W^{1118}$** | Biology Centre CAS<br>Czech Republic |
| **w-; Bl/Cyo; TM2/TM6B** | MPI-Jena, Germany |
| **w; $Or22ab^{GAL4}$** | Benton Lab, Switzerland |
| **w-; +/+; UAS-OR(w+)/UAS-OR(w+)** | BestGene Inc, USA |

**Table S18.** Origin of chemicals.

| Panel | Compound | Provider | Purity |
|---|---|---|---|
| 1 | *n*-eicosane | Merck | 99% |
| 1 | *n*-docosane | Merck | 99% |
| 1 | 1-octadecanol | Merck | 99% |
| 1 | (3*Z*)-dodecen-1-ol | IOCB | in house synthesis, see supplementary methods, GC purity 95% |
| 1 | (3*Z*,6*Z*)-dodecadien-1-ol | IOCB | in house synthesis, see supplementary methods, GC purity 94% |
| 1 | (3*Z*,6*Z*,8*E*)-dodecatrien-1-ol | IOCB | in house synthesis, see supplementary methods, GC purity 96% |
| 1 | dodec-3-yn-1-ol | IOCB | in house synthesis, see supplementary methods,GC purity 93% |
| 1 | *δ*-cadinene | Chemenu | 95% |
| 1 | (3*R*,6*E*)-nerolidol | IOCB | in house synthesis (Havlíčková et al. 2019), 96% |
| 1 | neocembrene | IOCB | purified from natural resource (Sillam-Dussès et al., 2005), GC purity 96% |
| 1 | geranylgeraniol | Merck | 98% |
| 2 | myrcene | Sigma Aldrich | ≥90% |
| 2 | eucalyptol | Sigma Aldrich | 99% |
| 2 | (+)-3-carene | Sigma Aldrich | 90% |
| 2 | (+)-limonene | Thermo-Fisher | 97% |
| 2 | (+)-α-pinene | Sigma Aldrich | 98% |
| 2 | p-cymene | Sigma Aldrich | 99% |
| 2 | γ-terpinene | Sigma Aldrich | 97% |
| 2 | (-)-β-caryophyllene | Sigma Aldrich | 98% |
| 2 | (+)-longifolene | Phyto Lab | ≥90% |
| 2 | terpinolene | Sigma Aldrich | ≥85% |
| 2 | toluene | VWR Chemicals | 99% |
| 2 | (2*E*)-hexenal | Thermo-Fisher | 98% |
| 2 | sabinene | Chemenu | ≥95% |
| 2 | pentyl acetate | Sigma Aldrich | ≥99% |
| 2 | ethyl acetate | VWR Chemicals | 99% |
| 2 | γ-nonalactone | Sigma Aldrich | 98% |
| 2 | heptanal | Thermo-Fisher | 98% |
| 2 | isoamyl propionate | Sigma Aldrich | ≥98% |
| 2 | ethanol | VWR Chemicals | ≥99% |
| 2 | (2*E*)-hexen-1-ol | Sigma Aldrich | 96% |
| 3 | *β*-bisabolene | Thermo-Fisher | 96% |
| 3 | (*E*)-myrcenol | FytoFarm | 95% |
| 3 | (-)-trans-pinocarveol | Sigma Aldrich | ≥96% |
| 3 | isoamyl acetate | Sigma Aldrich | ≥99% |
| 3 | 2-phenylethyl acetate | J&K Scientific Ltd. | ≥98% |
| 3 | α-camphorene | Synergy Ltd | ≥90% |
| 3 | (*E*)-β-ocimene | TRC Canada | 98% |
| 3 | nonan-1-ol | Thermo-Fisher | 95% |
| 3 | octan-1-ol | Honey well | 99% |
| 3 | benzyl alcohol | Thermo-Fisher | 99% |
| 3 | p-cresol | Sigma Aldrich | ≥99% |
| 3 | cryptone | Chemenu | ≥95% |
| 3 | (+)-cis-carveol | BOC Sciences | 95% |
| 3 | methyl jasmonate | Sigma Aldrich | ≥98% |
| 3 | eucarvone | MuseChem | ≥95% |
| 3 | ethyl (2*E*,4*Z*)-decadienoate | Sigma Aldrich | 95% |
| 4 | (-)-verbenone | Merck | 94% |
| 4 | acetophenone | Sigma Aldrich | 99% |
| 4 | 2-phenylethanol | Acros organics | 99% |
| 4 | (±)-myrtenol | Sigma Aldrich | 95% |
| 4 | geranyl acetone | Sigma Aldrich | ≥97% |
| 4 | oct-1-en-3-ol | Thermo-Fisher | 98% |
| 4 | 4-vinylanisole | Sigma Aldrich | 97% |
| 4 | 4-ethylguaiacol | Sigma Aldrich | ≥98% |
| 4 | hexan-1-ol | Sigma Aldrich | 99% |
| 4 | oct-1-en-3-ol | Sigma Aldrich | ≥97% |
| 4 | styrene | Thermo-Fisher | 99% |
| 4 | 2,3-dihydrobenzofuran | Thermo-Fisher | 99% |
| 4 | 2-methylbutan-1-ol | J&K Scientific Ltd. | 98% |
| 4 | 2-methylbutyl acetate | Sigma Aldrich | 99% |
| 4 | 4-methylanisole | Sigma Aldrich | 97% |
| 4 | linalool | Thermo-Fisher | 97% |
| 4 | methyleugenol | Sigma Aldrich | 98% |
| 4 | (±)-camphor | Sigma Aldrich | ≥95% |
| 4 | benzaldehyde | Sigma Aldrich | ≥99% |
| 4 | isoamyl alcohol | VWR Life Science | ≥98% |

### SUPPLEMENTARY METHODS

#### Organic synthesis

Unless noted otherwise, all reactions were carried out under argon in oven-dried glassware. Solvents were distilled from drying agents as indicated and transferred under nitrogen: THF (Na/benzophenone), toluene (Na/benzophenone). All starting materials were used as purchased (Sigma Aldrich, Combi-Blocks), unless otherwise indicated. Chromatography was performed using Fluka silica gel 60 (0.040 - 0.063 mm). For TLC analysis, F254 – coated aluminum sheets were used. The spots were detected both in UV and by the solution of $Ce(SO_4)_2{\cdot}4H_2O$ (1%) and $H_3P(Mo_3O_{10})_4$ (2%) in 10% sulfuric acid. $^1$H- and $^{13}$C NMR spectra were recorded at 400 MHz and 100 MHz, respectively, with a Bruker 400 MHz instrument at 25 °C (the solvents are indicated in parentheses). Chemical shifts are reported in ppm relative to TMS. The residual solvent signals in the $^1$H and $^{13}$C NMR spectra were used as an internal reference ($CDCl_3$: δ = 7.26 for $^1$H and δ = 77.23 for $^{13}$C). The compounds were analyzed using the gas chromatograph TRACE 1310 (ThermoFisher Scientific, Waltham, MA, USA), equipped with a nonpolar Zebron ZB-5MS column (30 m × 0.25 mm × 0.25 µm film; Phenomenex, Torrance, CA, USA) connected to a ThermoFisher Scientific ISQ LT mass-selective detector (70eV ionization voltage, source temperature 200 °C, transferline heated to 260 °C). The column temperature was held at 50 °C for 1 min, gradually increased to 320 °C at 8 °C/min and then held at 320 °C for 20 min. Helium was used as carrier gas at a flow 1.2 mL/min. Split/splitless port was heated to 200 °C, and samples were injected in splitless mode with a purge time of 1 min.

Scheme 1.

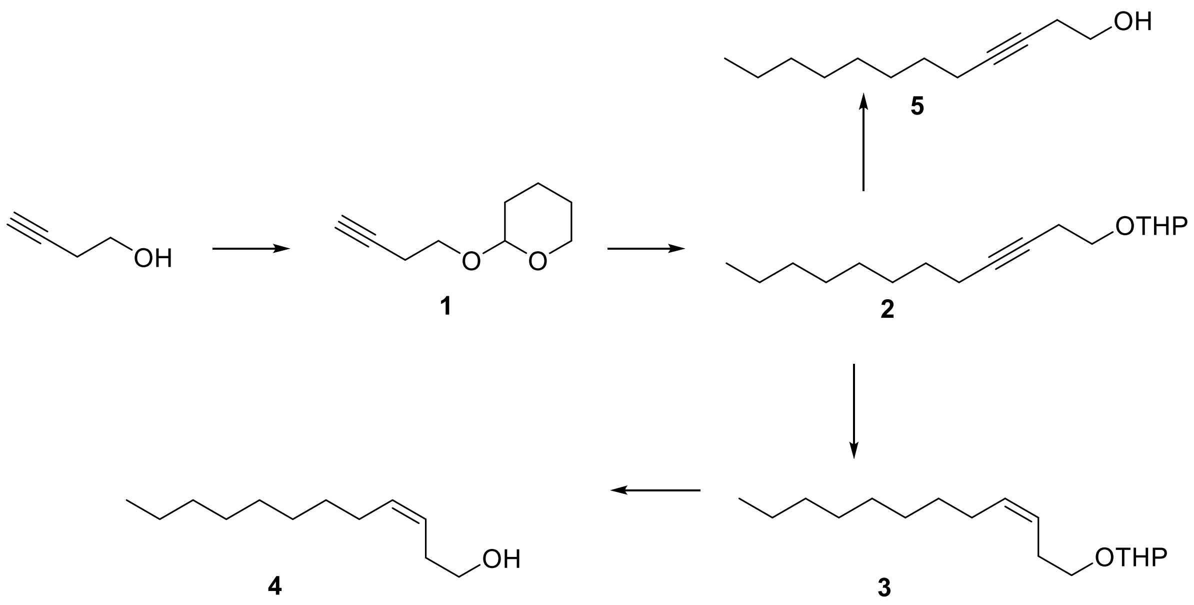

**2-(But-3-yn-1-yloxy)tetra-2H-hydropyran (Allegretti & Ferreira, 2011) (1)**

To a solution of 3-butyn-1-ol (3.0 g; 42.79 mmol) in DCM (100 mL) at 0 °C was added 3,4-dihydro-2*H*-pyran (3.78 g; 44.93 mmol) followed by addition of *p*-toluenesulfonic acid monohydrate (0.080 g; 0.427 mmol). After stirring at 0 °C for 5 min, the reaction was warmed to room temperature and stirred for 90 min. The reaction was quenched with saturated solution of $NaHCO_3$ (20 mL), and the layers were separated. The aqueous layer was extracted with DCM (50 mL), and the combined organic layers were dried over $MgSO_4$ and concentrated in vacuo. The residue was purified by a column chromatography (silica gel; cyclohexane/EtOAc 4:1) to give **4** as colorless oil (0.53 g, 64%). $^{1}$H NMR (401 MHz, $CDCl_3$) δ 4.67 (dd, *J* = 4.2, 2.9 Hz, 1H), 3.97 – 3.81 (m, 2H), 3.65 – 3.49 (m, 2H), 2.52 (td, *J* = 7.0, 2.7 Hz, 2H), 2.00 (t, *J* = 2.7 Hz, 1H), 1.93 – 1.78 (m, 1H), 1.78 – 1.69 (m, 1H), 1.69 – 1.46 (m, 4H). EI-MS (70 eV): *m/z* (%): 153 (1.4), 125 (2), 99 (9), 85 (100), 79 (9), 67 (21), 53 (33), 41 (22).

Scheme 2.

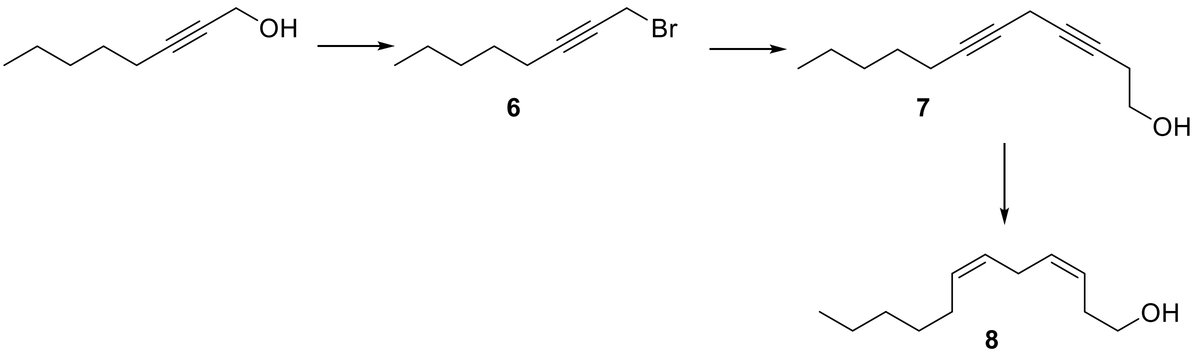

#### 1-Bromooct-2-yne (6)

A mixture of 2-octynol (2.0 g; 15.87 mmol), triphenylphosphine (4.57 g; 17.46 mmol), and tetrabromomethane (6.30 g; 19.04 mmol) in DCM (40 mL) was stirred at 0 °C for 2 hr and then it was diluted with cyclohexane and filtered. The filtrate was evaporated, and a column chromatography (silica gel; eluent cyclohexane) of the residue afforded 3.70 g (90%) of bromide **5** as a pale oil. $^{1}$H NMR (401 MHz, $CDCl_3$) δ 3.95 (t, *J* = 2.4 Hz, 2H), 2.25 (ddt, *J* = 7.2, 4.7, 2.4 Hz, 2H), 1.59 – 1.46 (m, 2H), 1.44 – 1.27 (m, 4H), 0.96 – 0.88 (m, 3H). $^{1}$H NMR data match published spectrum (Sigurjónsson & Haraldsson, 2024).

Scheme 3.

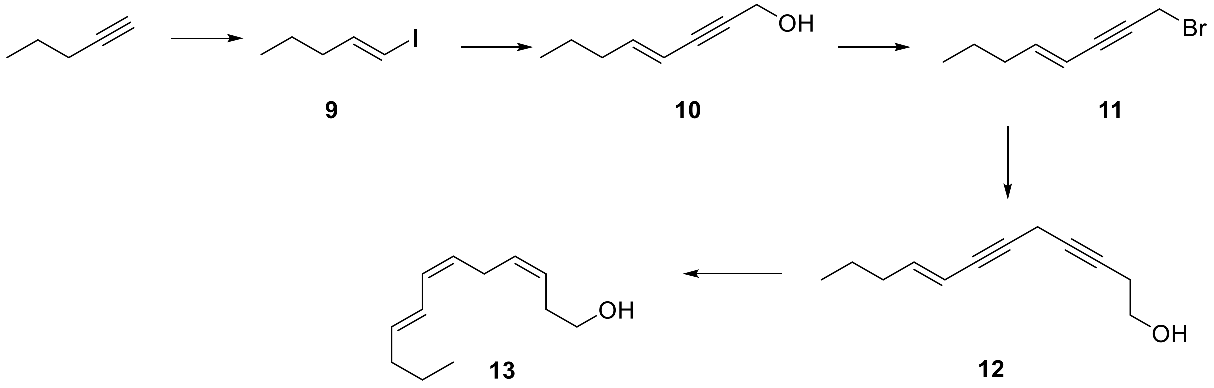


**(*E*)-1-iodopent-1-ene (Saran et al., 2018) (9)**

Diisobutylaluminum hydride in hexanes (1 M, 50 mL, 50 mmol) was added dropwise to a solution of pent-1-yne (3.7 g; 54 mmol) in hexanes cooled to -40 °C. The resulting mixture was stirred for 30 min, then slowly warmed to room temp over 3 hr, and was allowed to stir overnight. The mixture was warmed to ~50 °C for 4 hr, then cooled to -40 °C, and a solution of iodine (12.7 g; 50 mmol) in THF (20 mL) was added dropwise over 30 min. The resulting dark brown suspension was allowed to warm to room temperature with stirring overnight. The solution was cooled in an ice bath, and slowly quenched with dropwise addition of ice-cold diluted sulfuric acid (5 mL of 96% acid in 60 mL of water) with vigorous stirring. After stirring for 30 min, the layers were separated, and the organic layer was washed with dilute $NaHSO_3$ solution and brine and dried over $MgSO_4$. The resulting solution was filtered through a short pad of silica gel, rinsing with pentane. Obtained solution was concentrated by rotary evaporation without heating (rotavap was set to 100 Torr pressure). This procedure gave 12.24 g of crude alkenyl iodide **9** that was used in the next step. EI-MS (70 eV): *m/z* (%): 155 (4), 141 (3), 127 (5), 73 (69), 55 (100).

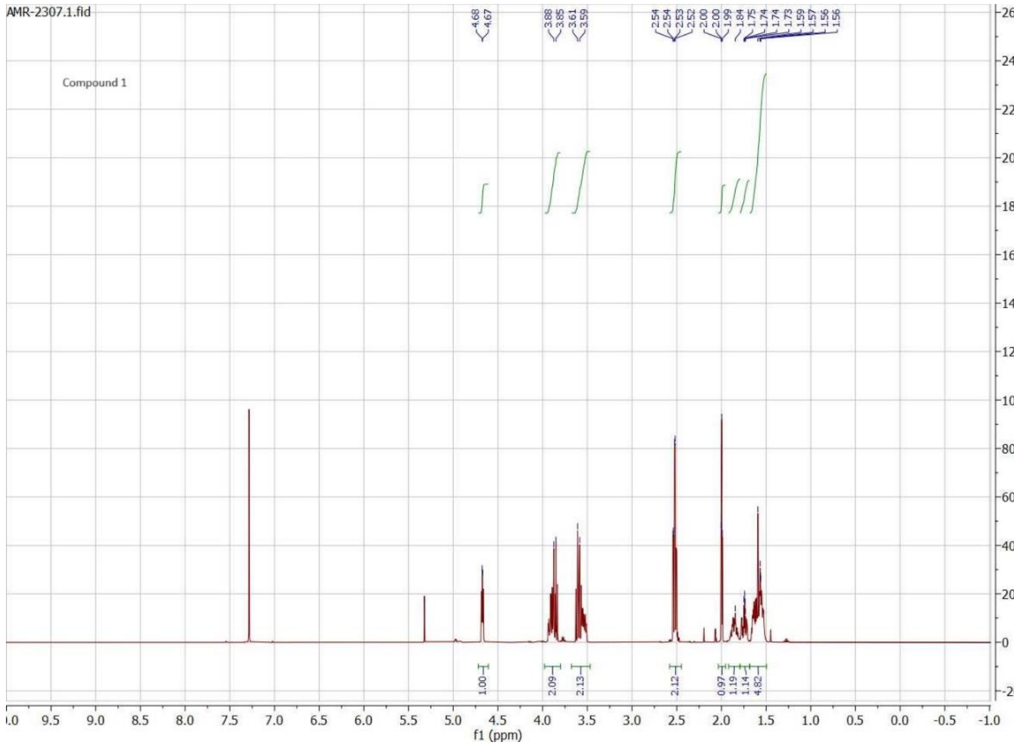


[1]H NMR spectrum of the compound 1.

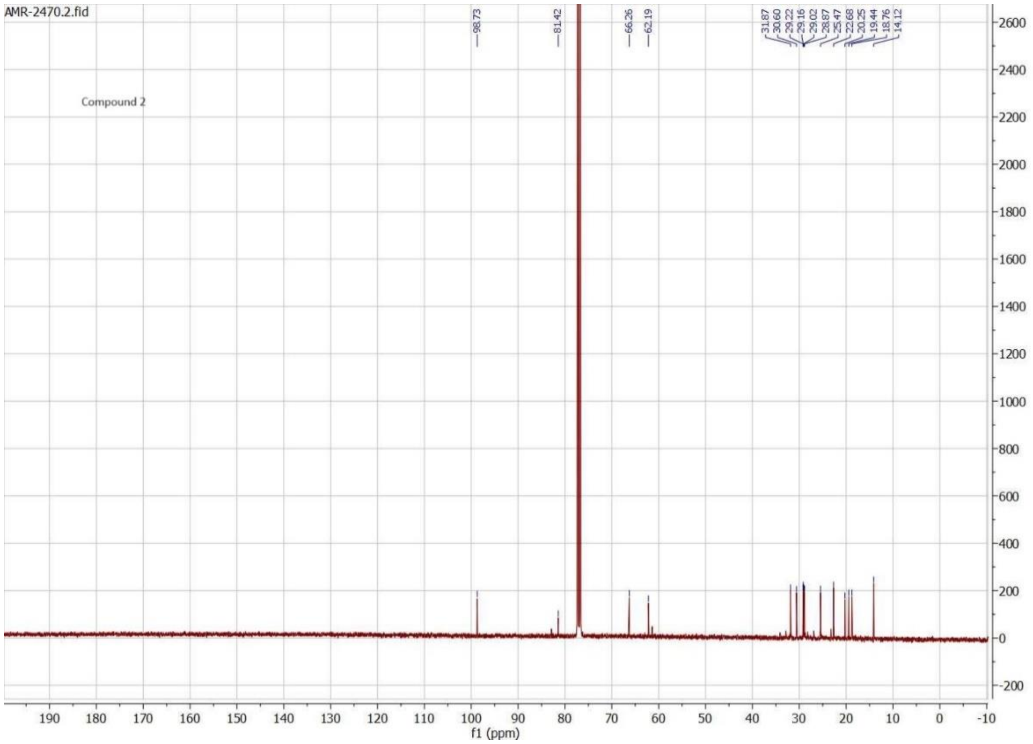


$^{13}$C NMR spectrum of the compound 2.

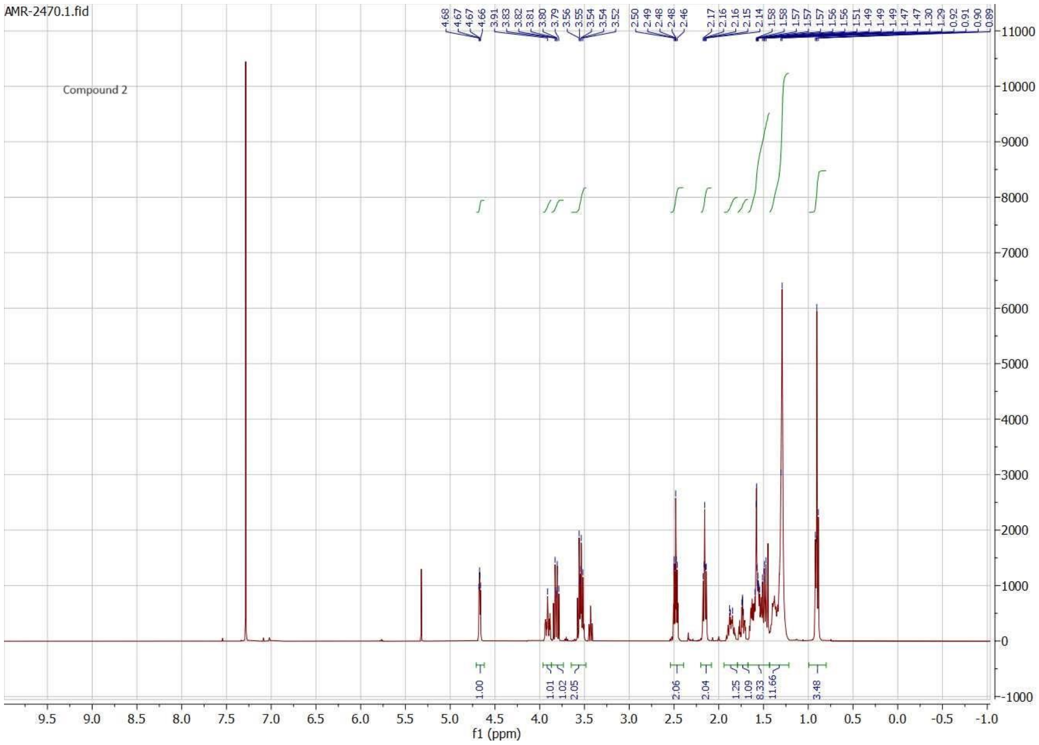


$^{1}$H NMR spectrum of the compound 2.

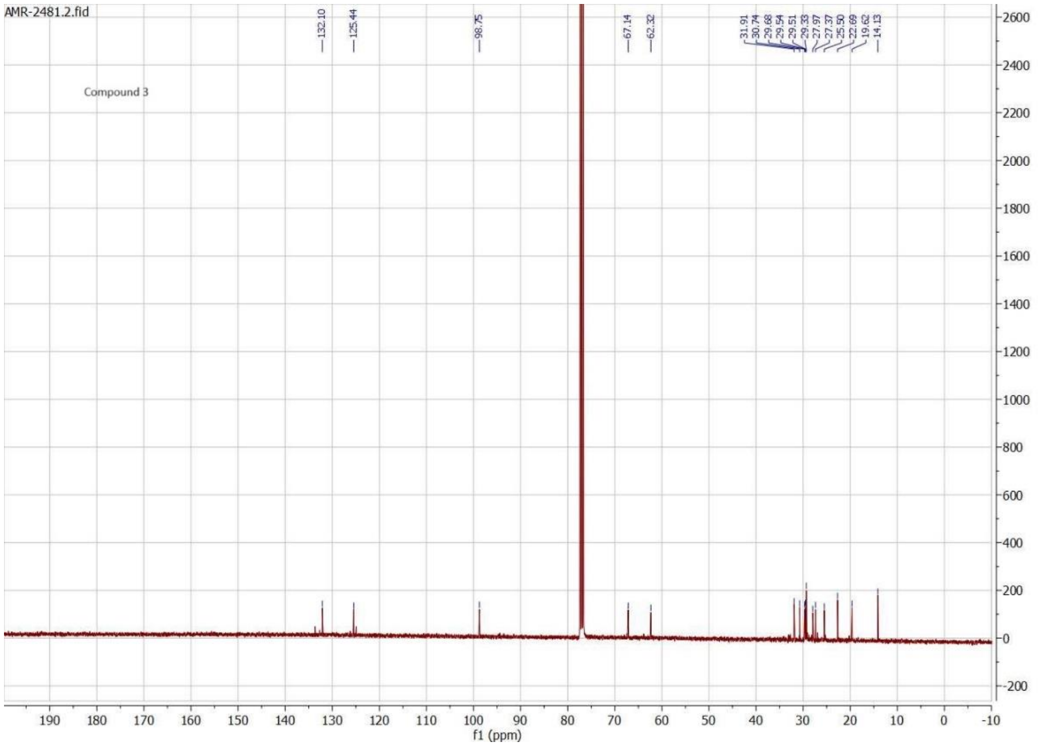


$^{13}C$ NMR spectrum of the compound 3.

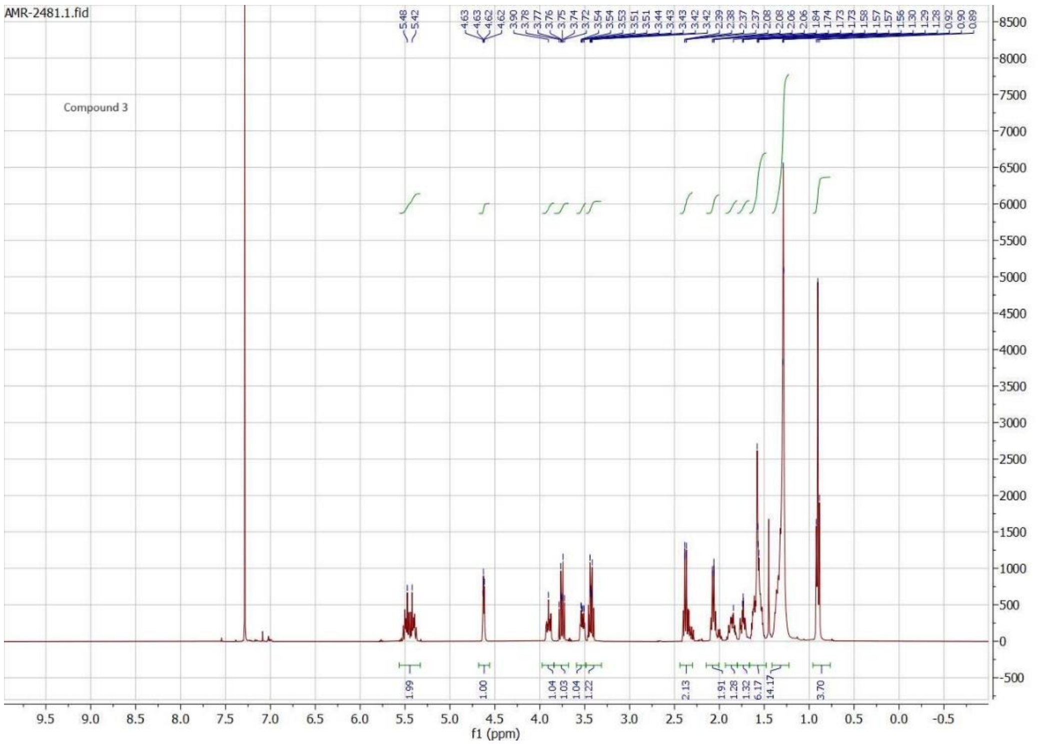


$^{1}H$ NMR spectrum of the compound 3.

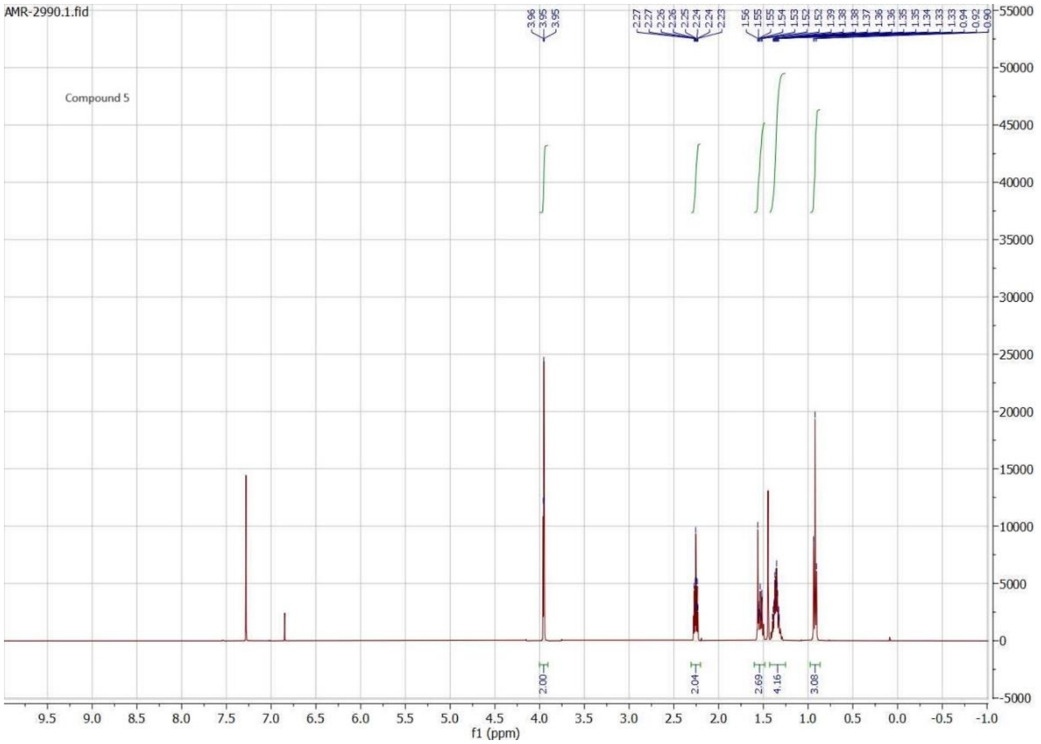


[1]H NMR spectrum of the compound 5.

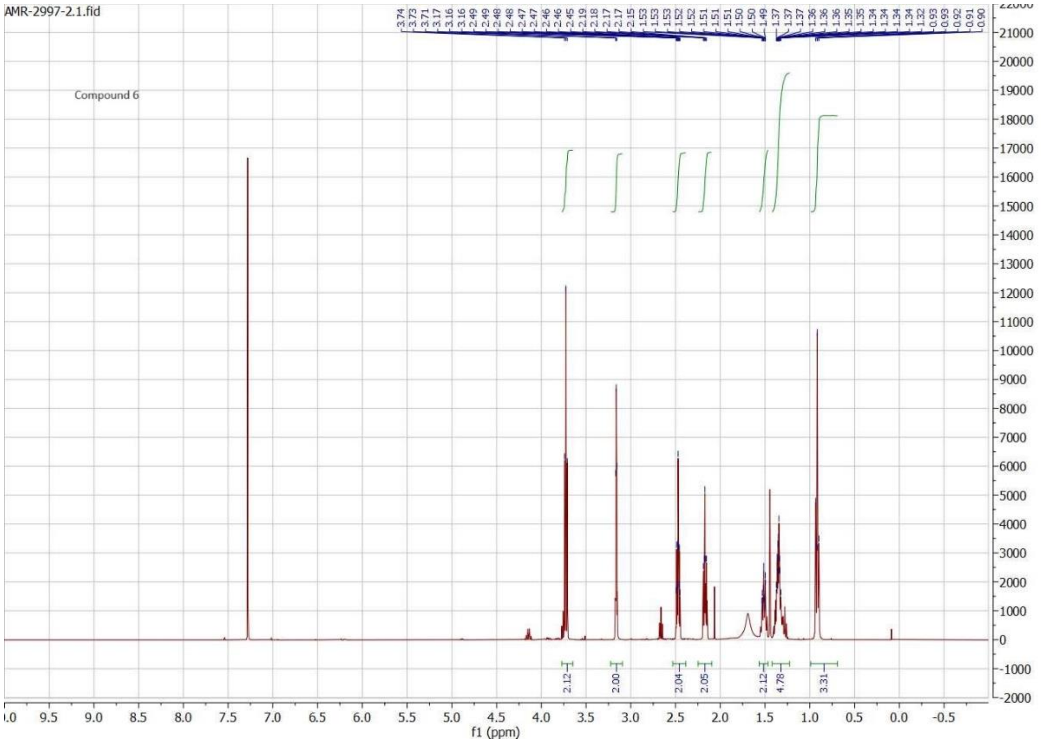


[1]H NMR spectrum of the compound 6.

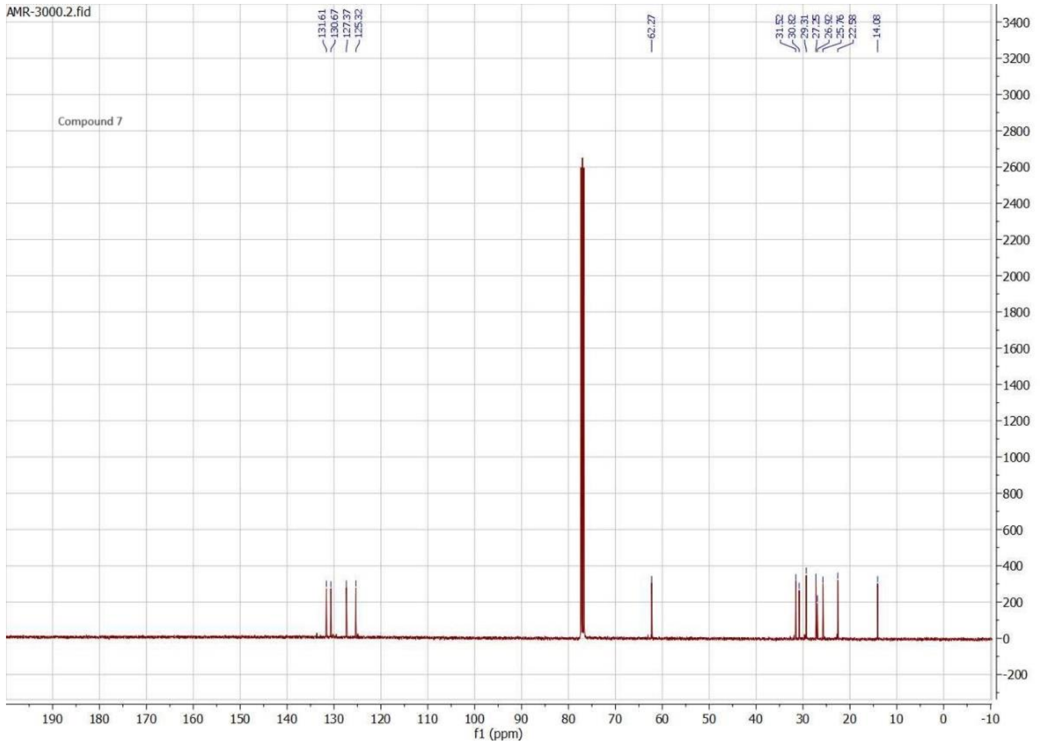


[13]C NMR spectrum of the compound 7.

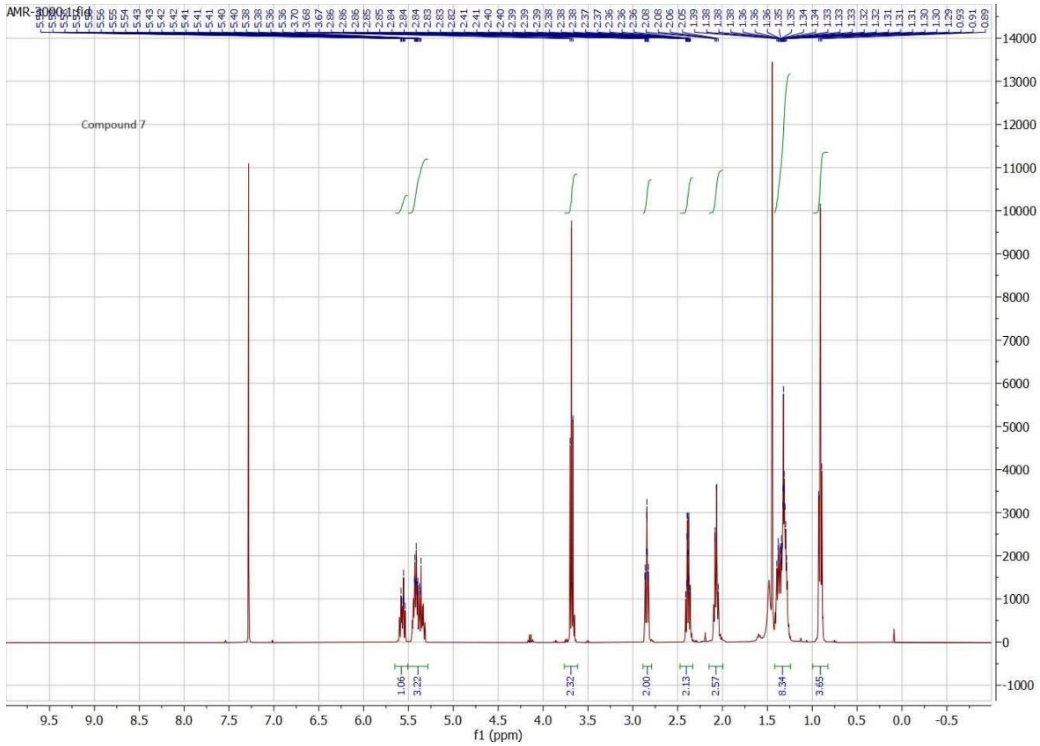


[1]H NMR spectrum of the compound 7.

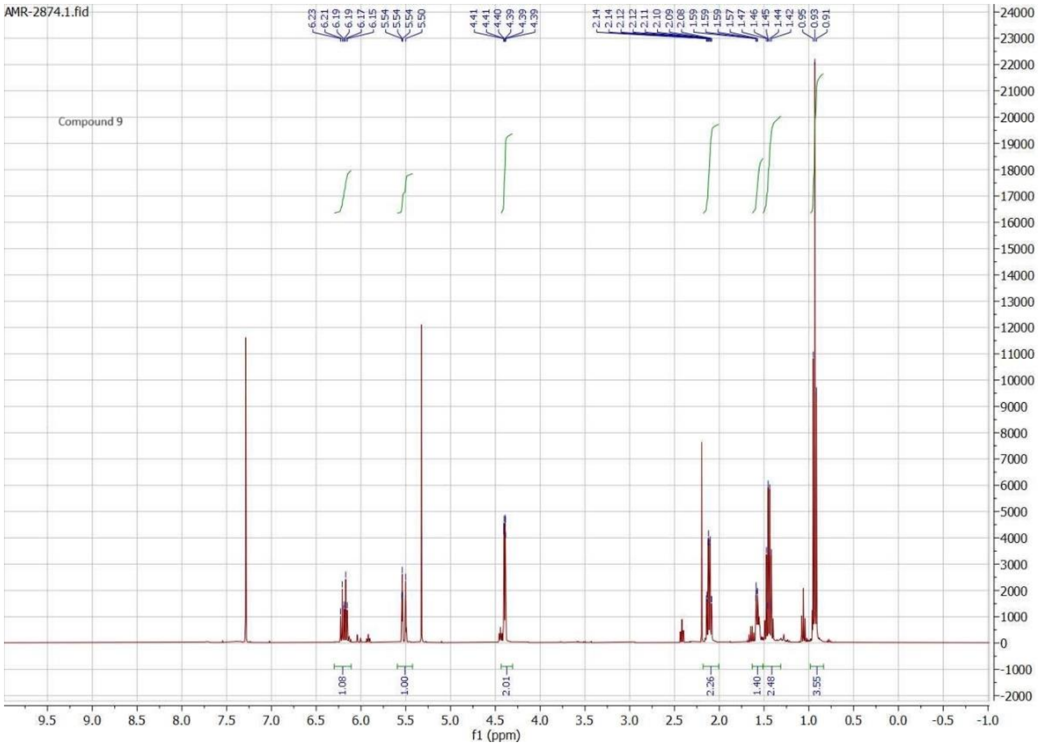


$^{1}$H NMR spectrum of the compound 9.

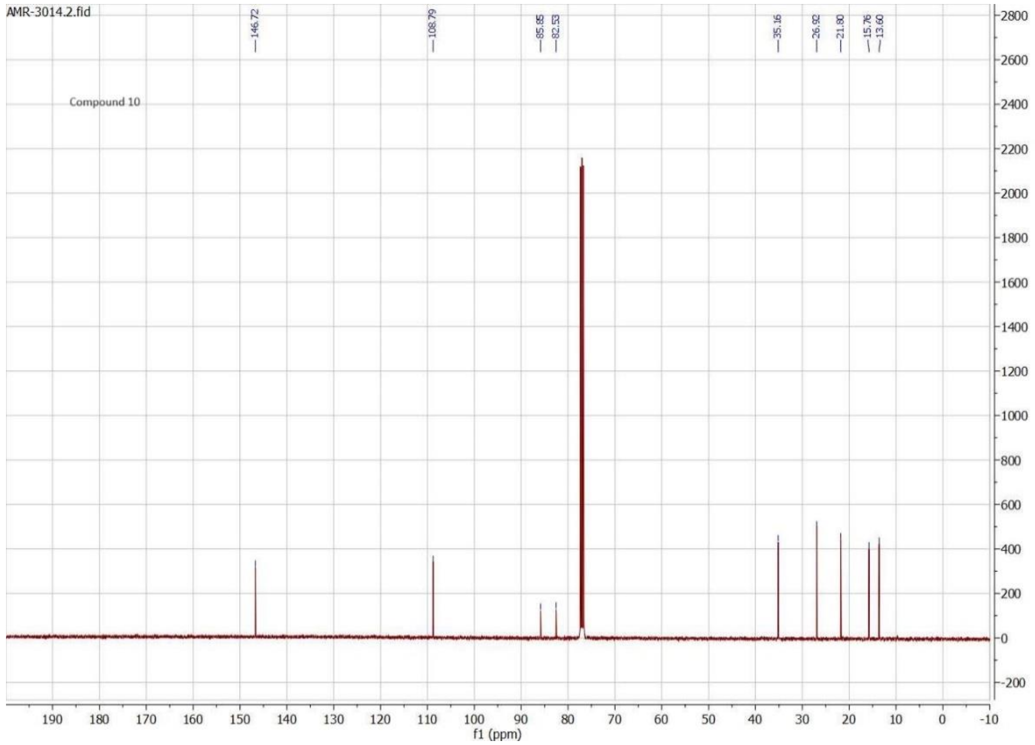


$^{13}C$ NMR spectrum of the compound 10.

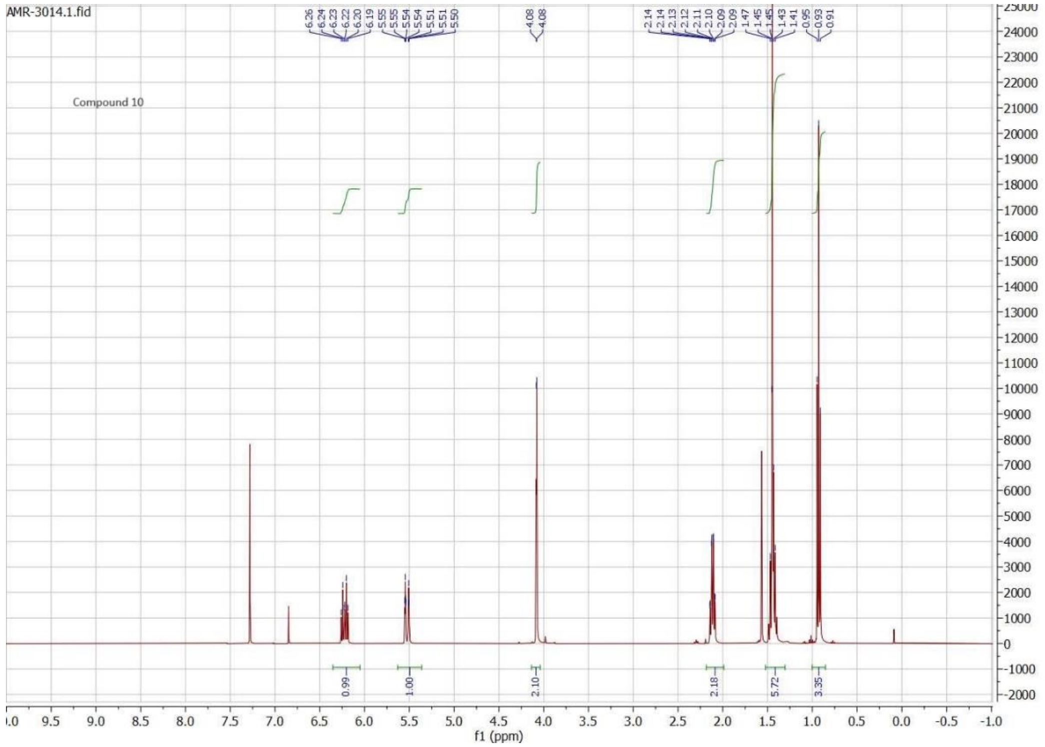


$^{1}H$ NMR spectrum of the compound 10.

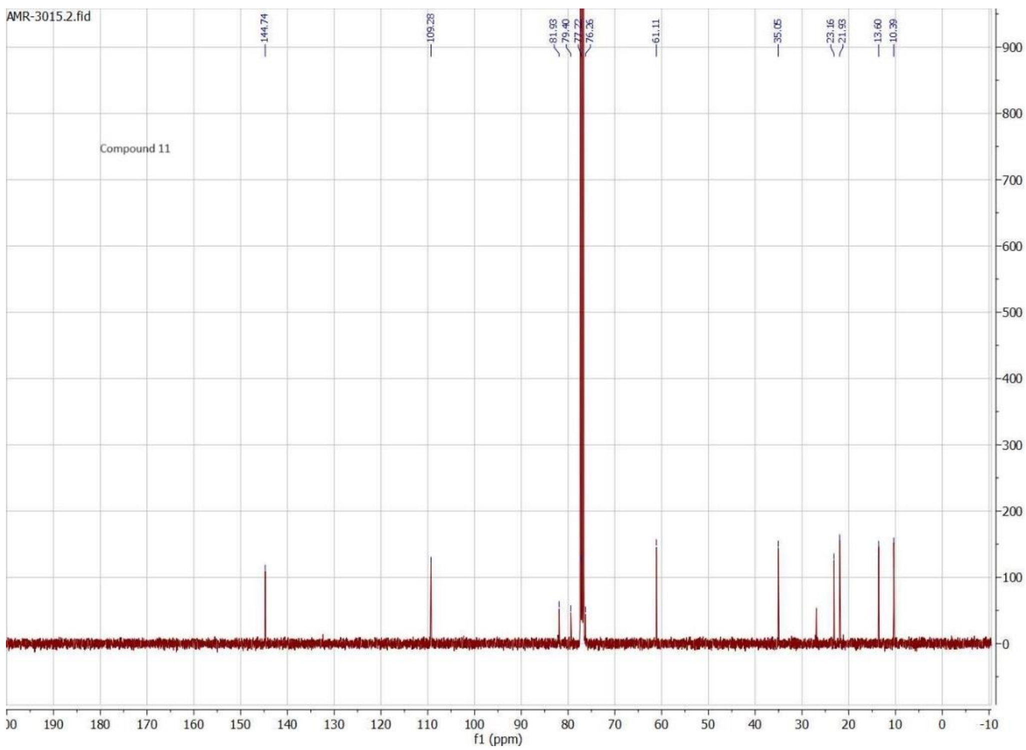


$^{13}C$ NMR spectrum of the compound 11.

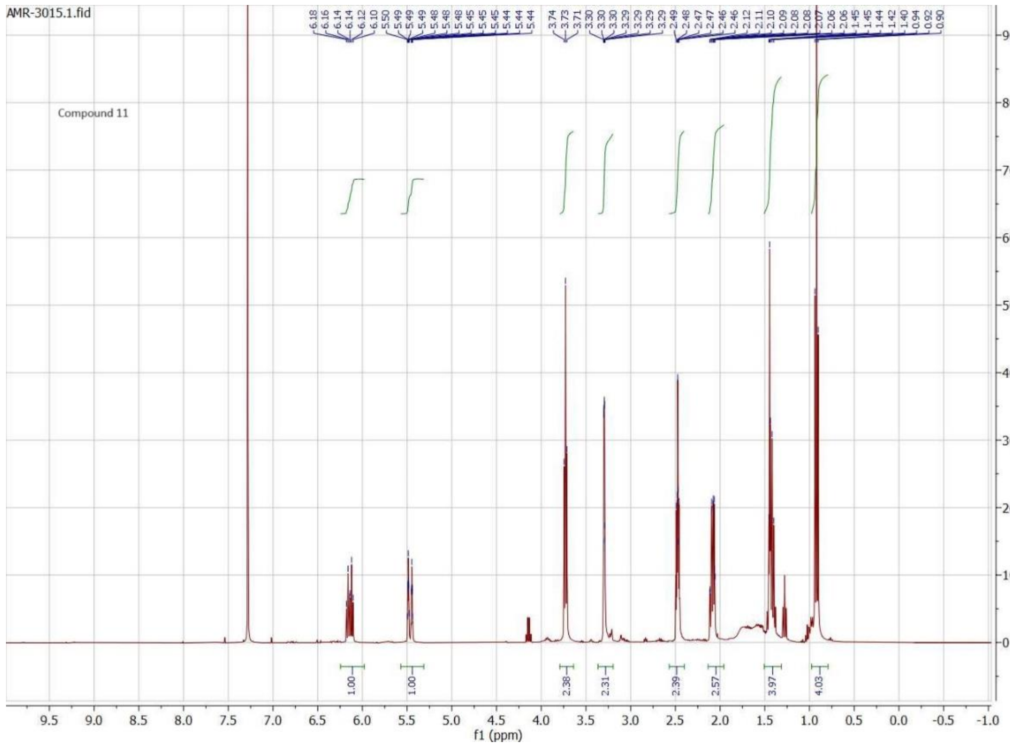


$^{1}H$ NMR spectrum of the compound 11.

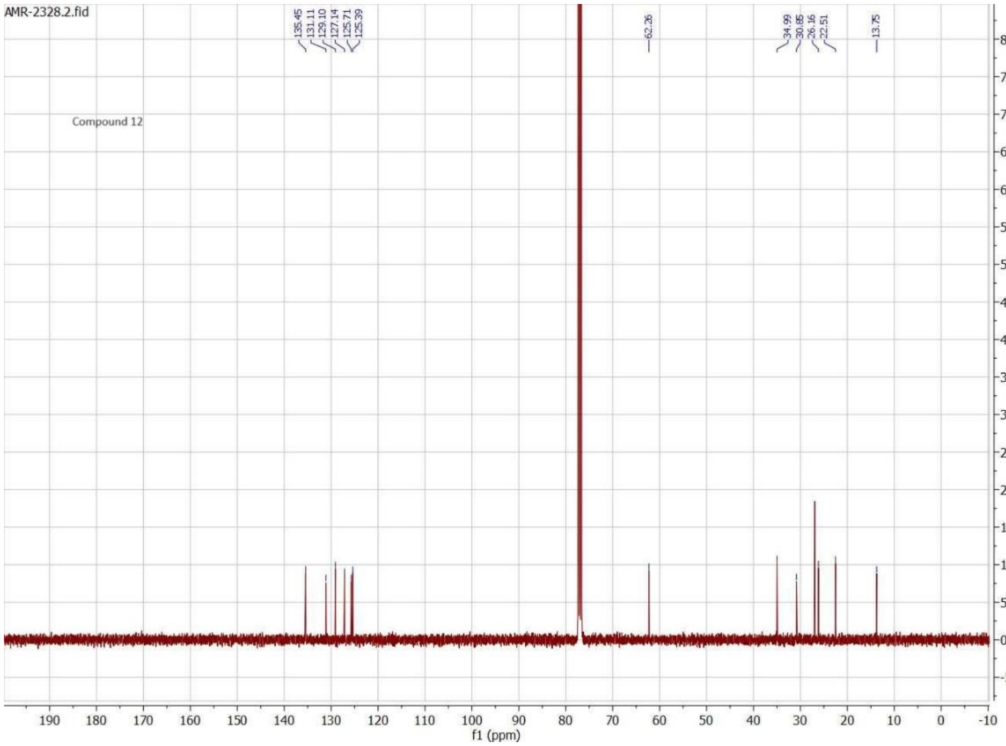


$^{13}$C NMR spectrum of the compound 12.

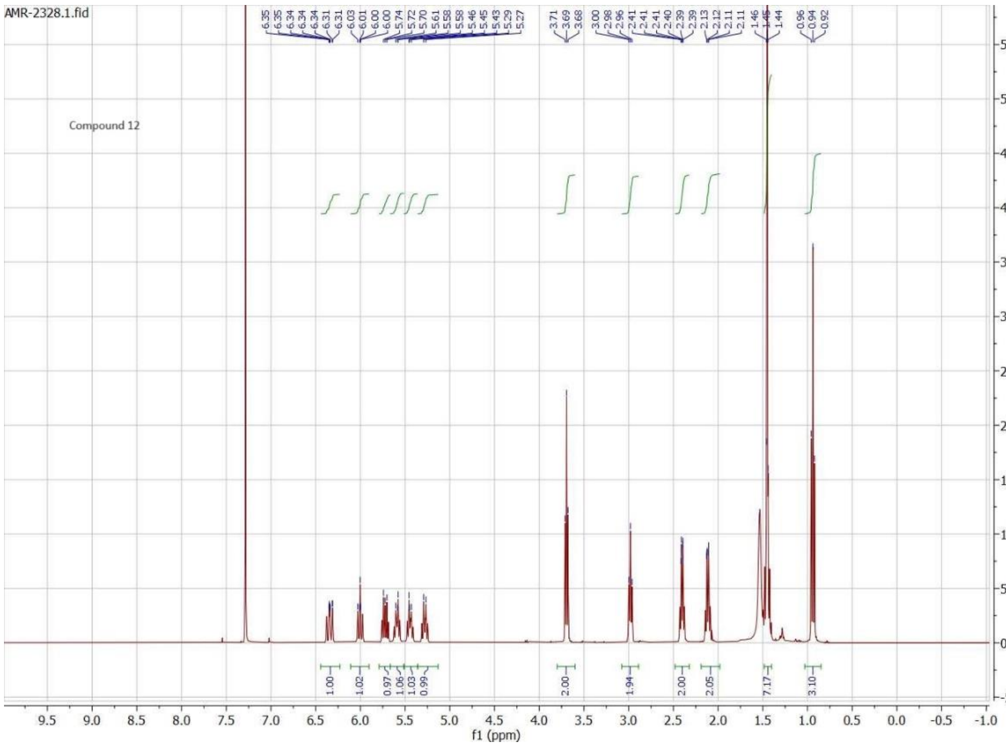


$^{1}$H NMR spectrum of the compound 12.
